# Decoding the Impact of Lipid Saturation on ER Signaling Networks, ERSU, UPR, and ERAD

**DOI:** 10.1101/2025.09.15.676312

**Authors:** Xia Li, Maho Niwa

## Abstract

The faithful inheritance of a functional endoplasmic reticulum (ER) in *Saccharomyces cerevisiae* is safeguarded by the ER Stress Surveillance (ERSU) checkpoint, which delays cytokinesis when ER homeostasis is perturbed. Under stress, ER transmission to the daughter cell is halted, while in parallel—but through independent pathways—the Unfolded Protein Response (UPR) restores ER function and ER-associated degradation (ERAD) eliminates misfolded proteins, ultimately allowing cell cycle re-entry. ER stress also transiently stimulates sphingolipid biosynthesis, with the intermediate phytosphingosine (PHS) acting as a key activator of ERSU. Yet, how broader lipid parameters—such as membrane composition, saturation, and fluidity—reshape ER quality control and, in particular, govern ER inheritance during division remains poorly understood. To address this, we employed a tightly controlled experimental system to selectively alter lipid saturation and phospholipid composition while monitoring ER inheritance within the framework of ER homeostasis maintained by UPR and ERAD. Strikingly, we found that perturbations in lipid balance exerted specific effects on ER inheritance that were distinct from their impact on UPR and ERAD. These findings reveal lipid homeostasis as a critical integrator of ER functional regulation, linking ERSU, UPR, and ERAD into a unified adaptive network that ensures robust ER transmission and cellular resilience under stress.

## Introduction

In eukaryotic cells, essential biochemical reactions are compartmentalized within membrane-bound organelles, ensuring spatial and regulatory control over diverse cellular processes. Among these, the endoplasmic reticulum (ER) stands out as one of the largest and most functionally diverse organelles. It serves as a central hub for the synthesis, folding, and maturation of proteins destined for secretion or integration into cellular membranes—collectively classified as secretory pathway proteins^1–7^. These proteins are assisted by molecular chaperones to ensure correct folding and undergo a series of tightly regulated post-translational modifications, including glycosylation, disulfide bond formation, and oligomerization, before being transported to the Golgi apparatus and beyond.

The ER is also the principal site of lipid biosynthesis, producing a broad spectrum of phospholipids, sterols, and lipid-derived molecules that are vital for maintaining the structural integrity and functional capacity of intracellular membranes. Beyond serving as the structural matrix of cellular compartments, lipids regulate membrane properties such as curvature, fluidity, and thickness—parameters that, in turn, influence membrane protein folding, insertion, and trafficking. Recent studies have drawn increasing attention to the intimate link between ER lipid composition and its functional capacity: imbalances in lipid saturation or phospholipid availability can impair protein folding, alter membrane protein dynamics, and disrupt ER morphology^8^. Elevated levels of saturated fatty acids (SFAs) increase membrane rigidity, disrupt lipid–protein interactions, and trigger defects in vesicle trafficking, calcium signaling, and protein quality control^9–11^. Conversely, imbalances in specific lipid species—such as phosphatidylinositol depletion—can compromise ER function by affecting lipid-dependent signaling pathways, membrane fluidity, and organelle communication^12, 13^. Importantly, the ER is also a major calcium storage organelle, and its ability to coordinate calcium homeostasis is intimately tied to lipid composition, further influencing cell signaling and metabolic outcomes.

Given its diverse physiological roles, the ER must continuously adapt to changes in the cellular environment while maintaining homeostasis. Perturbations such as protein misfolding or lipid imbalance initiate ER stress and activate a suite of adaptive quality control mechanisms that function cooperatively to restore ER function and protect against cellular damage^14, 15^. Furthermore, these events are coordinated with cell cycle, upon adjustment of the ER inheritance. In the budding yeast *Saccharomyces cerevisiae*, the ER is compartmentalized into two major domains: the perinuclear ER (pnER), which envelopes the nucleus, and the cortical ER (cER), which forms a dynamic network underneath the plasma membrane. As the ER cannot be synthesized de novo during cell division, it must be faithfully inherited from the mother cell. ER inheritance begins with the formation of an initial ER tubule (IET) that emerges from the pnER and extends toward the bud site in early cell cycle stages^16, 17^. This tubule later expands and establishes a functional cER network in the daughter cell. Accurate ER inheritance is critical, as its failure results in compromised ER function in daughter cells and can lead to cell death^18–20^.

To preserve ER integrity and function during stress, eukaryotic cells rely on three major partially overlapping stress response pathways: the ER Stress Surveillance (ERSU) pathway, ER-associated degradation (ERAD), and the unfolded protein response (UPR). These pathways detect and respond to distinct yet often interrelated forms of ER dysfunction, forming a coordinated and robust quality control network^3, 21–29^. The ERSU pathway functions as a stress-activated cell cycle checkpoint that halts ER inheritance during mitosis, thereby ensuring that only functional ER is transmitted from mother to daughter cells. When ER stress is detected, ERSU activation blocks cER inheritance and cytokinesis, preserving daughter cell viability by preventing the inheritance of damaged ER^20, 30, 31^. ERAD, in contrast, maintains proteostasis by identifying and retro-translocating misfolded proteins from the ER to the cytosol, where they are degraded by the ubiquitin-proteasome system^23, 32^. Substrate recognition in ERAD is categorized by the location of the misfolded domain—luminal (ERAD-L), membrane-associated (ERAD-M), or cytosolic (ERAD-C)—and handled by distinct ubiquitin ligase complexes (e.g., Hrd1, Doa10) in conjunction with the Cdc48 ATPase^33, 34^. Molecular mechanisms of ERAD and the impact of lipid changes, specifically increased lipid saturation or depletion of essential phospholipids—in modulating ERAD efficiency and substrate selectivity have been described. The third key pathway, the UPR, responds to ER stress by triggering widespread transcriptional and translational reprogramming to restore ER folding capacity and expand the ER network. In yeast, UPR engagement is initiated through the ER-resident sensor Ire1, which activates the unconventional splicing of *HAC1* mRNA, producing the Hac1 transcription factor that upregulates genes involved in protein folding, ERAD, and lipid biosynthesis^35^. Notably, emerging evidence shows that lipid abnormalities—such as increased membrane saturation or impaired phospholipid synthesis—can directly activate the UPR independently of misfolded protein accumulation upon integrating both proteotoxic and lipotoxic signals to gauge ER homeostasis via Ire1^8, 24^.

While the detail mechanisms by which different forms of lipid imbalance modulate UPR intensity, duration, or cross-talk with other stress response pathways remain largely unknown. While extensive work has defined the independent roles and mechanisms of ERSU, ERAD, and UPR in responding to classical protein misfolding stress, the integrated response of these pathways under conditions of membrane lipid imbalance remains elusive. Specifically, how perturbations in membrane composition—such as alterations in lipid saturation, changes in phospholipid flux, or defects in lipid biosynthetic pathways—impact ER inheritance, protein degradation, and stress signaling thresholds remains.

To address these knowledge gaps, we employed a multifaceted approach to manipulate ER lipid composition and assess its effects on ER quality control. First, we used a previously established experimental system^36^ to gradually increase ER membrane lipid saturation, that allowed us to examine how changes in membrane fluidity affect ER functions. In parallel, we assessed ER homeostasis under inositol depletion, a condition that disrupts phosphatidylinositol synthesis and alters the cellular phospholipid pool, thereby imposing a distinct form of lipid stress without necessarily increasing saturation. We also utilized targeted genetic mutants defective in key steps of lipid biosynthesis to isolate the contributions of specific lipid classes.

Using these complementary strategies, we analyzed how shifts in lipid homeostasis affect each of the major ER stress response pathways—ERSU, ERAD, and UPR. Our findings reveal that increased membrane saturation and phospholipid imbalance impair cER inheritance, sensitize cells to UPR activation, and selectively disrupt ERAD processing of membrane-associated substrates. These results establish ER membrane composition not merely as a passive backdrop but as a dynamic and critical regulator of ER quality control. More broadly, our study highlights the central role of lipid-protein interactions in shaping cellular responses to stress, advancing our understanding of how membrane biology integrates with proteostasis and cell cycle progression.

## Results

### ER Lipid Saturation via OLE1 Regulates Cortical ER Inheritance at Late Stages

To investigate whether alterations in ER lipid saturation influence the unfolded protein response (UPR), the ER surveillance (ERSU) pathway, and ER-associated degradation (ERAD), we employed a previously established system in *Saccharomyces cerevisiae* that modulates lipid saturation through promoter variants controlling *OLE1* expression²⁹ (**Fig. *S1A***). *OLE1* encodes the sole fatty acid desaturase in yeast, introducing double bonds into saturated fatty acids (FA) which results in changes of the membrane fluidity. Gradual reduction of *OLE1* expression was achieved by promoter mutagenesis, generating four yeast strains (SFA1-SFA4) with progressively increased membrane phospholipid (PL) saturation and a decreasing phosphatidylethanolamine (PE)/phosphatidylcholine (PC) ratio. Mass spectrometry analysis from the previous study confirmed that SFA1 featured a wild-type PL composition, which was followed by SFA2 and SFA3 with moderate increases, and finally, SFA4 displayed the most pronounced increase in saturated FA, which was characterized by a marked rise in fully saturated phospholipid species and shortened acyl chain length (decreasing from ∼C16–C18 species toward shorter chains)^36^. In addition, the PE/PC ratio decreased across the SFA strains, reflecting broader remodeling of ER membrane lipid composition.^36^

To visualize lipid saturation levels within different ER sub-regions, particularly the inner nuclear membrane (INM) and perinuclear ER (pnER), we applied Mga2-GFP–based lipid saturation (LipSat) reporters based on Mga2-GFP^37^. Mga2 is an ER-resident transcription factor whose GFP-tagged form localizes to both cortical ER (cER) and pnER^31^ (**Fig. *S1B–C***). Under elevated lipid saturation, Mga2 undergoes proteolytic cleavage within its transmembrane domain, releasing a nuclear-targeted fragment that accumulates in the nucleoplasm^32,33^. Conversely, under reduced saturation, Mga2 remains membrane-bound without cleavage. Quantification of GFP signals in the nucleoplasm versus ER membranes (cER/pnER) thus provides a rapid readout of the relative distribution of saturated versus unsaturated lipids in ER membrane or INM^30^. In wild-type control cells (MNY3371), ∼25% of ER LipSat sensor GFP fluorescence localized to the membrane fraction (unsaturated), while ∼75% appeared in the nucleus (saturated) (**Fig. *S1D***). Supplementing cultures with oleic acid (C18:1), which moderately reduces lipid saturation, led to a slight decrease in Mga2 cleavage and nuclear GFP accumulation. Linoleic acid (C18:2), which more strongly reduces saturation, further decreased Mga2 cleavage and favored membrane localization of the sensor (**Fig. *S1D***). Comparable patterns were observed for the INM LipSat sensor GFP fluorescence (**Fig. *S1E***). These outcomes mirrored prior studies^30^ and validated the reporter’s performance in our system.

Applying this ER LipSat reporter to the SFA strains, we observed that SFA1 behaved similarly to wild type (∼60% unsaturated, ∼40% saturated), while SFA2, SFA3, and particularly SFA4 showed progressively higher Mga2 nuclear signals, demonstrating elevated lipid saturation (**Fig. 1A–B**). We next asked how these lipid changes influenced cortical ER (cER) inheritance, a process we assayed using Pho88- GFP, an integrated ER reporter^21^. As previously described, ER inheritance proceeds in a sequential manner in budding yeast^16,34^: during early cell cycle stages (class I), an initial ER tubule (IET) extends from the pnER toward the bud neck, enters the daughter cell, and anchors at the polarisome at the bud tip. This is followed by lateral spreading along the plasma membrane to establish the cortical ER beneath the daughter cortex (class II–III cells). Consistent with earlier reports, tunicamycin-induced ER stress abolished cER inheritance in many class I cells (**Fig. 1C–D**). When cells encounter stress much later (class III), cells completed division, but cER inheritance was blocked in the next cycle^21,35^.

**Figure 1:**
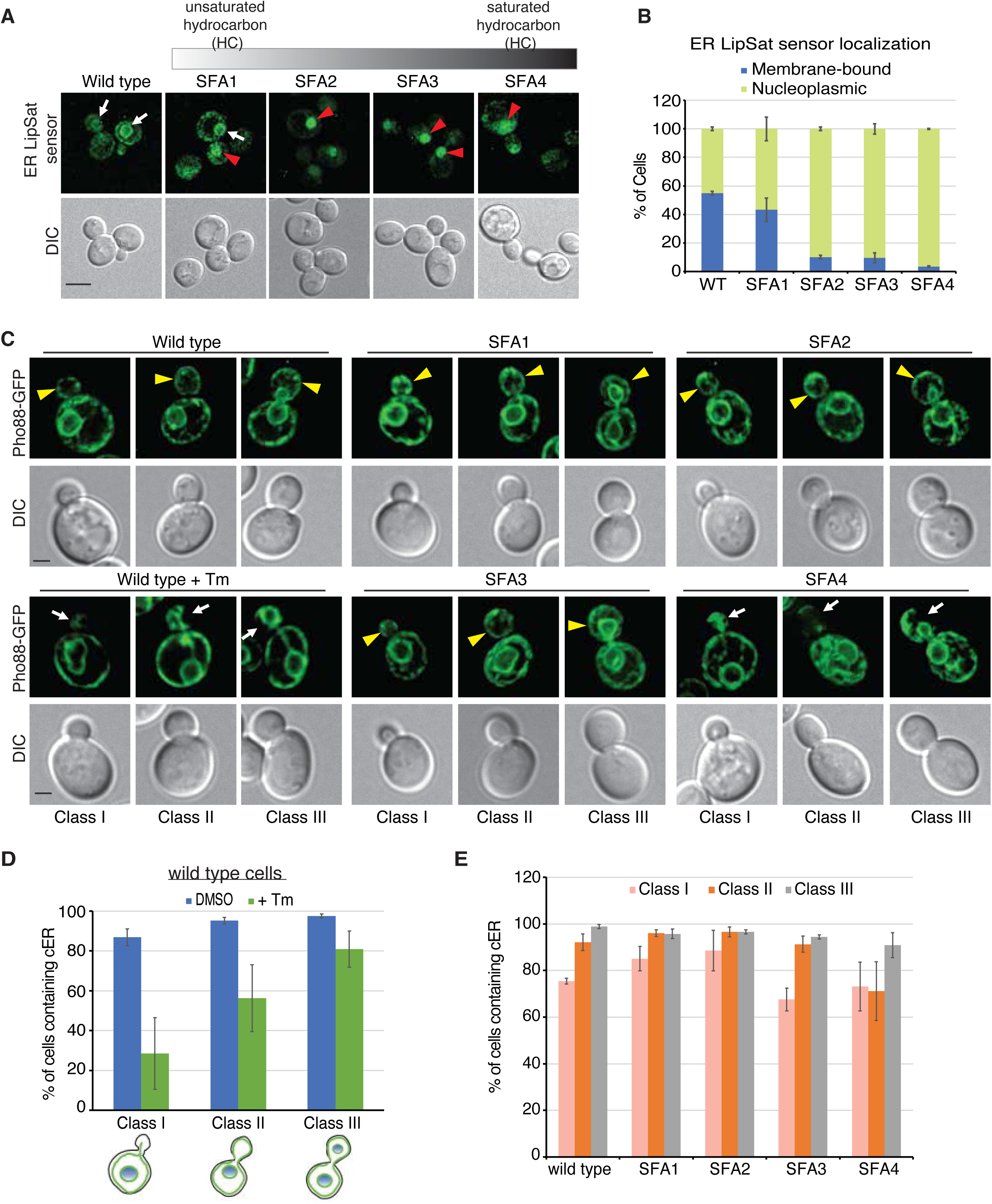
Lipid Saturation level has effects on ER inheritance. (**A–B**) Analysis of ER lipid saturation levels in SFA1–SFA4 strains. (**A**) Representative fluorescence images showing localization of the ER LipSat sensor in wild type (MNY3414, W303 background) and SFA1-SFA4 strains. Cells were grown in standard SC medium containing 2% glucose. A grayscale bar above the images illustrates gradual changes in levels of saturated fatty acids. White arrows mark membrane-associated ER LipSat sensor, while red arrowheads indicate nucleoplasmic localization. (B) Quantification of membrane-bound versus nucleoplasmic ER LipSat sensor localization. Fluorescence distribution was measured from >100 cells per strain. Scale bar, 5 μm. (**C**) Cortical ER (cER) inheritance defects in SFA4. ER was visualized with a well-established ER marker Pho88-GFP. Representative Live-cell images of wild-type and SFA1–SFA4 strains expressing Pho88-GFP are shown. For comparison, wild-type cells treated with 1 μg/mL tunicamycin (Tm) for 90 min were included to induce ER stress. Yellow arrowheads indicate normal cER in daughter cells, while white arrows point to defective cER. Cells were classified into categories I–III as previously described^30^. Scale bar, 2 μm. (**D**)-(**E**) Quantification of the cER inheritance in class I, II and III for WT (**D**) and SFA1-SFA4 cells (**E**). Over 100 cells per class were analyzed, with data representing the average of three independent experiments. Error bars denote standard deviation (SD).

In the SFA background, cER inheritance defects correlated with the degree of lipid saturation (**Fig. 1C, E**). SFA1 and SFA2 resembled wild type. In contrast, SFA3 class I cells, and SFA4 class I–II cells, displayed significantly reduced cER inheritance, indicating that early stages of inheritance are particularly sensitive to increased lipid saturation. In SFA4, the reduction was strongest in class II cells, although still less severe than the inheritance blocks triggered by tunicamycin (**Fig. 1D–E, *S2A***). SFA3 and SFA4 thus exhibited unique inheritance profiles, with SFA4 showing the clearest impairment, especially in class II daughters, distinguishing it from the relatively intact inheritance seen in WT, SFA1, and SFA2.

To investigate whether early steps of ER inheritance were disrupted in SFA4, we analyzed IET initiation and orientation in synchronized class I or II cells. Cells were arrested at G2/M with hydroxyurea and released into G1, capturing populations enriched for incipient bud formation (**Fig. 2A–B**). Using calcofluor white (CFW) staining to visualize bud scars, we monitored IET orientation relative to the bud neck. Operationally, IETs were defined as tubular extensions emerging from the pnER toward the bud. Quantitative imaging revealed no significant differences between WT and SFA4 in IET frequency, timing, length, or intensity, nor in the proportion of tubules reaching into the bud (**Fig. 2B–F**). Thus, early ER targeting events remained largely intact in SFA4.

**Figure 2:**
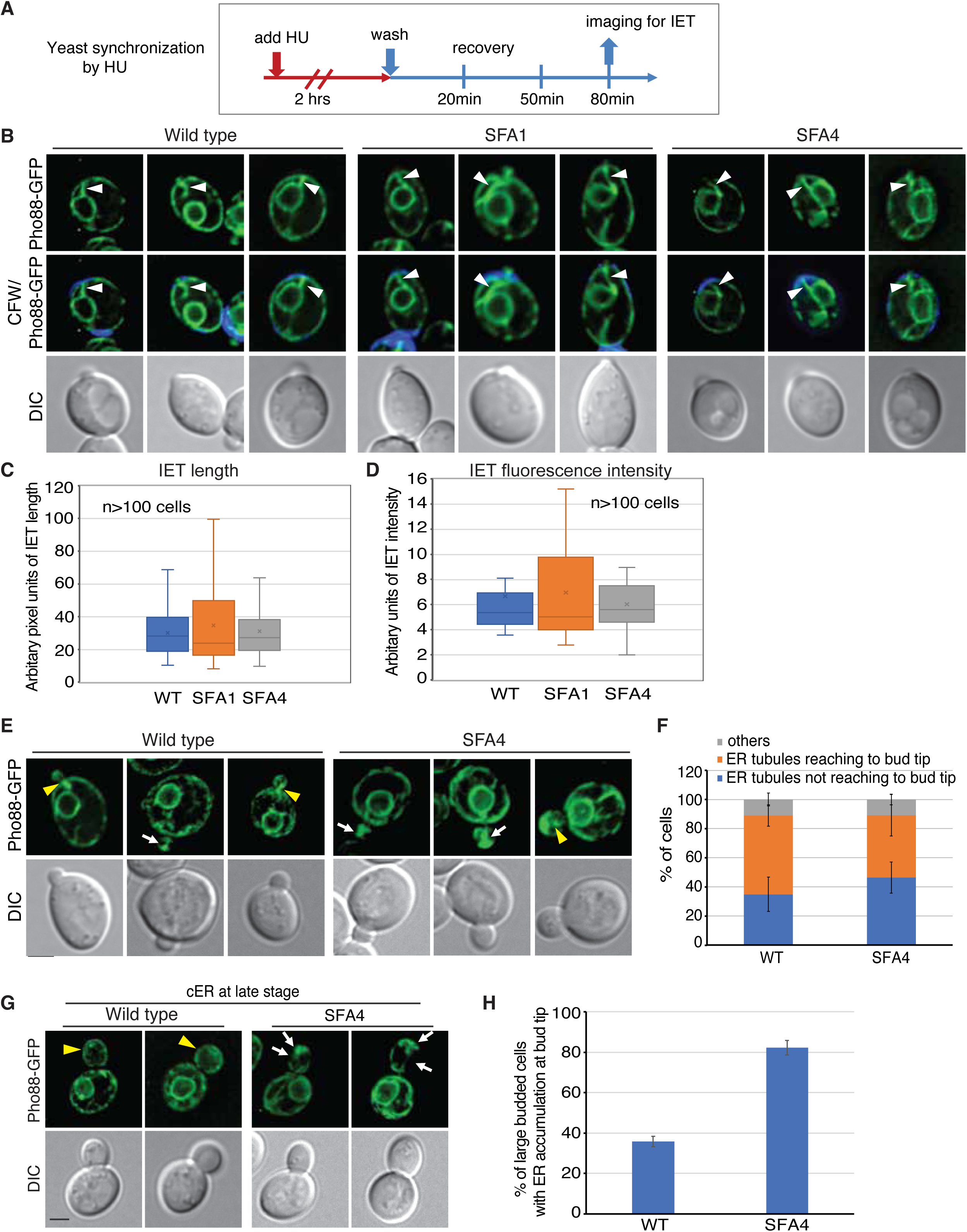
Effects of Lipid Saturation level on different stages of cortical ER (cER) inheritance. (**A**) Experimental diagram for hydroxyurea (HU) synchronization release into cell cycle. Yeast cells were synchronized in S-phase by treatment with 100 mM hydroxyurea (HU) for 2 hours. After HU treatment, cells were washed to remove HU and allowed to recover in fresh medium. Samples were collected at 20, 50, and 80 min post-wash to monitor progression through the cell cycle. Imaging for initial ER tubules (IET) was performed at 80 minutes after HU release. (**B**) The formation of initial ER tubules (IET) and their entry into the new bud in early stages of cER inheritance are not affected in SFA4 cells. Calcofluor White (CFW) staining marks cytokinetic remnants (CRMs). Three representative cells are shown for each indicated strain, with images from the GFP channel, merged GFP and DAPI channels, and DIC. Arrowheads indicate an IET extending from the perinuclear ER (pnER) in the mother cell toward the newly forming bud. Scale bar, 5 µm. (**C–D**) Quantification of IET tubule length and fluorescence intensity across different strains. Measurements were performed using ImageJ as described in Materials and Methods. Over 100 cells were analyzed per strain. (**E**) Representative images showing initial ER tubules (IET) extending into the bud at different stages of cER inheritance in wild-type and SFA4 cells. Yellow arrowheads indicate cells with an IET extending a half way into the daughter cell but not reach to the bud tip, while white arrows highlight IET extension reaching fully to bud tip. Scale bar, 5 µm. (**F**) Quantification of cells exhibiting different ER tubule positions. Depending on the inheritance stage, ER tubules either dynamically extend to the bud tip or fail to do so. (**G**) cER in large buds during the late stages of cell cycle does not extend all the way through the cortex of the daughter cell in SFA4 cells. Yellow arrowheads indicate normal cER distribution in wild-type cells, while white arrows highlight ER accumulation at the bud tip and a lack of cER around the bud cortex in SFA4 cells. Scale bar, 2 µm. (**H**) Quantification of cER accumulation at the bud tip in large-budded wild-type and SFA4 cells.

Striking defects emerged instead in **class II cells**. Compared to ∼30% of WT daughters, nearly 80% of SFA4 class II buds accumulated ER markers abnormally at the bud tip, with little or no cortical ER along the bud cortex (**Fig. 2G–H**). Although these inheritance defects were milder than those observed under acute ER stress, SFA4 uniquely exhibited stronger impairment in class II than in class I or III cells. Taken together, these findings indicate that excess ER lipid saturation does not disturb the earliest targeting events of ER inheritance but selectively disrupts later stages, particularly the maturation and distribution of cortical ER in class II daughter cells.

### Differential Effects of ER Lipid Saturation on ERAD Substrate Degradation

Recognizing the essential role of phospholipid homeostasis in ER function and cellular stress adaptation, we next explored how the saturation state of ER membrane lipids modulates the efficiency of ER-associated degradation (ERAD). Specifically, we quantified the half-lives of distinct ERAD substrates by monitoring their steady-state protein levels over time following cycloheximide (CHX)–mediated inhibition of protein synthesis—a well-established method to determine protein degradation dynamics (**Fig. 3**). By arresting new protein synthesis with CHX, we could precisely track the decrease in substrate abundance as a direct readout of degradation, allowing robust determination of protein half-life and stability under varying lipid saturation conditions. Using this cycloheximide chase assay, we probed the fate of three well-established ERAD substrates: Hmg2-GFP—an integral membrane protein processed via the ERAD-M pathway (mutation in the membrane domain), CPY*-HA—a mutated soluble luminal protein (G255R in carboxypeptidase Y) cleared by the ERAD-L pathway (luminal domain misfolding), and Ste6-166-3xHA- GFP (Ste6*-HA-GFP)—a mutant membrane protein with a misfolded cytosolic domain, handled by the ERAD-C pathway (cytosolic domain misfolding) (**Fig. 3A-F**). Strikingly, we found that Hmg2-GFP degradation was markedly accelerated in both SFA1 and SFA4 mutants compared to wild-type, with SFA1 exhibiting the most dramatic effect (**Fig. 3A-B**). The degradation of Ste6*-HA-GFP was also enhanced in both mutants but was most pronounced in SFA4 (**Fig. 3C-D**). In contrast, the clearance of CPY*-HA remained essentially unchanged across wild-type, SFA1, and SFA4 backgrounds (**Fig. 3E-F**).

**Figure 3:**
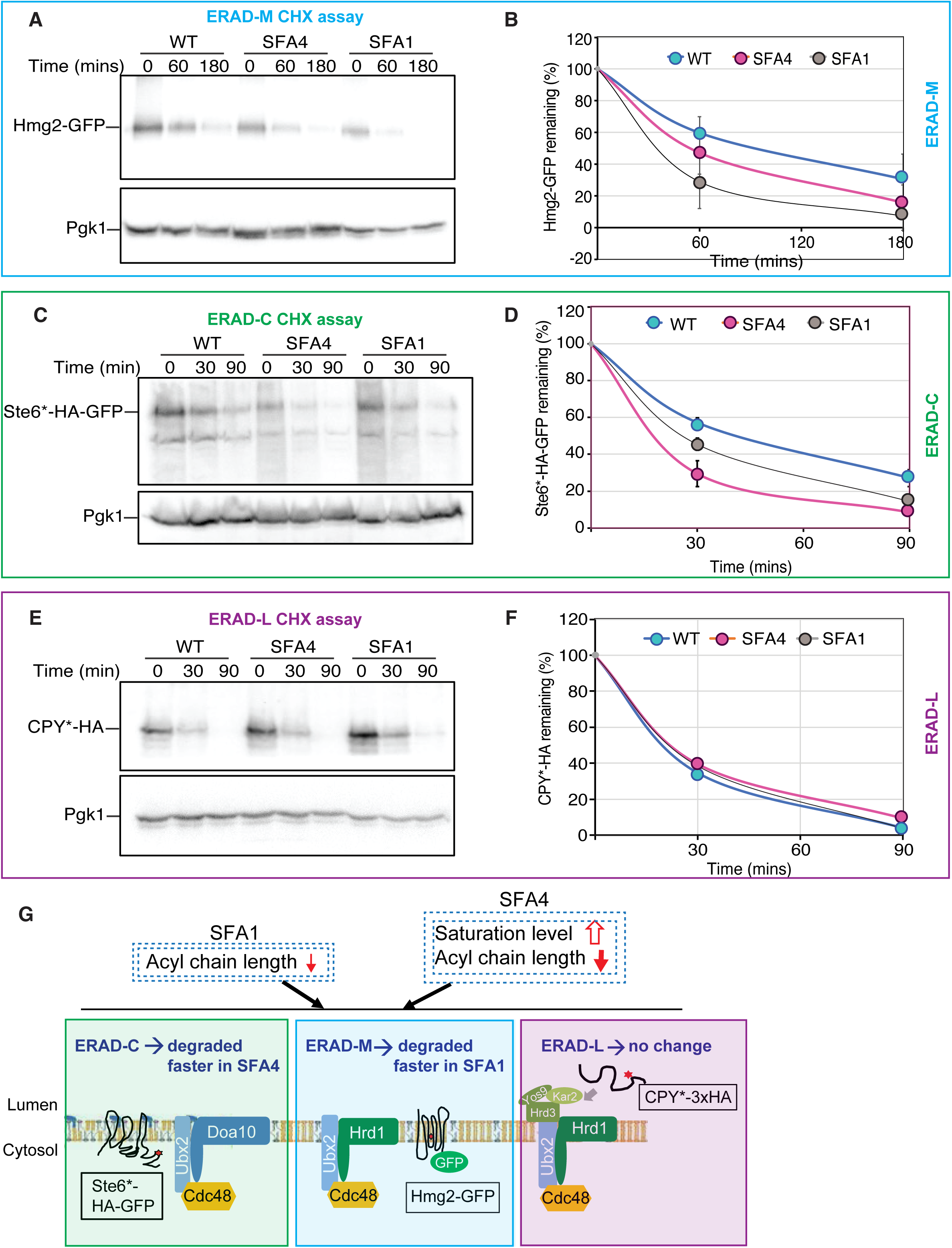
Effects of Lipid Saturation level on ERAD. (**A**)-(**E**) A cycloheximide (CHX) chase of Hmg2-GFP (A-B), CPY*-HA (C and D), and Ste6*-HA-GFP (E and F) expressed in Wild-type, lipid saturation mutants, SFA1 and SFA4. Upon CHX treatment, equal numbers of cells were collected to prepare total cell extracts for SDS-PAGE, followed by western blot analysis to detect ERAD membrane (ERAD-M) substrate Hmg2-GFP, ERAD luminal substrate (ERAD-L) CPY*-HA, and ERAD cytosolic substrate (ERAD-C), Ste6*-HA-GFP. Pgk1 was used as a loading control. The mean percentage of each ERAD substrate remaining in the cells for three biological replicates is plotted. Error bars represent the standard deviation (SD). (**G**) Schematic summary of ERAD substrate degradation in SFA1 or SFA4 yeast strains. In SFA1 yeast cells, membrane lipids display wild-type like saturation with slightly shortened acyl chain length. In contrast, SFA4 cells exhibit both markedly reduced acyl chain length and increased saturation of lipid membrane. These changes in membrane composition selectively enhance the degradation of ERAD-C and ERAD-M substrates–Ste6*-HA-GFP and Hmg2-GFP, respectively– while having no effect on the degradation of the ERAD-L substrate CPY*-3xHA. ERAD-C and ERAD-M substrates are recognized by the Doa10 and Hrd1 ubiquitin ligase complexes, respectively, and extracted by the Cdc48-Ubx2 complex. ERAD-L substrate degradation depends on the Hrd1-Hrd3 complex and the ER luminal chaperones Kar2 and Yos9.

These results reveal a specificity: membrane lipid saturation selectively accelerates the turnover of ERAD substrates with transmembrane or cytosolic misfolded domains, while leaving the degradation rate of soluble luminal proteins largely unaltered. Notably, SFA1 cells—enriched in unsaturated lipids— had a particularly strong impact on the stability of Hmg2-GFP, whereas SFA4 cells—with increased membrane saturation—most significantly reduced the half-life of Ste6*-HA-GFP (**Fig. 3G**). Together, our findings uncover distinct and substrate-specific influences of ER membrane lipid saturation on ERAD efficiency, underscoring the lipid environment as a powerful modulator of protein quality control within the ER.

### Lipid Saturation Levels of the ER Membrane Drive Distinct UPR Activation Dynamics

To determine whether increased ER membrane lipid saturation influences UPR activity, we monitored UPR induction using a UPRE-GFP reporter in wild-type (WT), SFA1, and SFA4 yeast cells. Cells were transformed with a well-established reporter plasmid expressing GFP under the control of a UPR transcriptional element (UPRE), and GFP fluorescence was quantified using ImageJ. As expected, treatment with tunicamycin (Tm; 1 µg/mL for 1.5 hours) led to an approximately two-fold increase in GFP fluorescence in WT cells, consistent with previous findings^24^. SFA1 cells treated with Tm exhibited GFP levels similar to those of Tm-treated WT cells (**Fig. 4A-B**).

**Figure 4:**
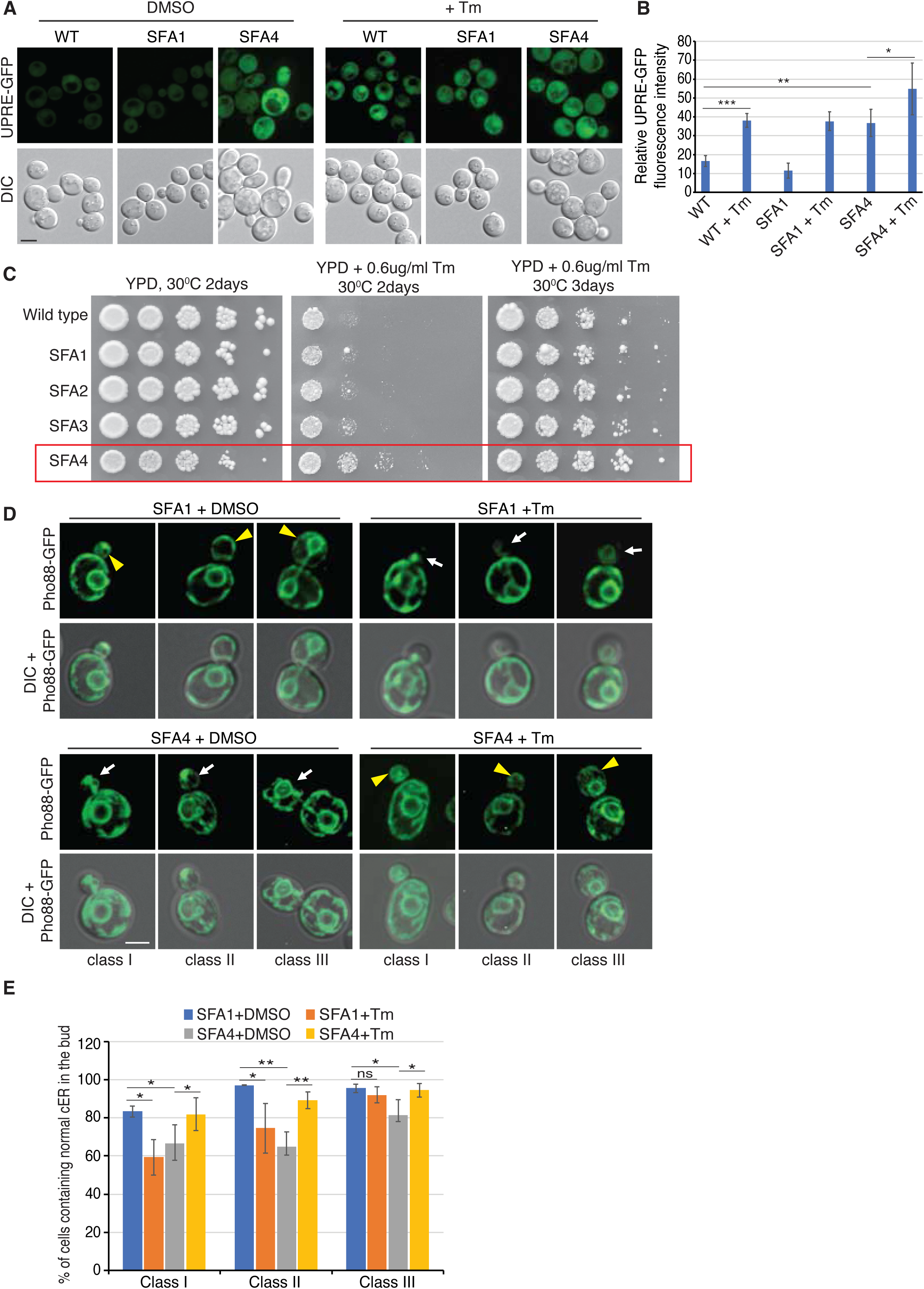
Effects of lipid saturation level on UPR pathways. (**A**) Expression of GFP in yeast cells carrying the UPRE-GFP reporter, where the unfolded protein response element (UPRE) was fused to GFP and expression of GFP reports levels of UPR activation by ER stress. Wild-type (WT), SFA1 or SFA4 yeast cells harboring the UPRE-GFP plasmid were incubated with control DMSO or 1 µg/ml Tm for 90 minutes for examining the ER stress activation levels. Representative images for each condition are shown. Scale bar, 5 µm. (**B**) Quantitating over 300 cells carrying UPRE-GFP, in either WT, SFA1 or SFA4 cells, with or without Tm treatment. GFP expression in SFA4 cells even prior to Tm treatment shows that SFA4 cells induce ER stress and activate UPR. Mean fluorescence intensity with standard error of the mean is shown. A one-tailed *t-test* was used to determine the significance of the difference between WT and SFA4 cells or between SFA4 cells with and without Tm treatment. p < 0.05, *; *p < 0.01*, **; p < 0.002, ***. (**C**) Growth phenotypes of SFA1 - SFA4 yeast strains with or without Tm. Serial dilutions of overnight cultures from the indicated strains were plated on YPD plates or YPD plates containing 0.6 µg/ml Tm and grown at 30°C for 2–3 days. The red box highlights the improved growth of the SFA4 strain on YPD plates containing 0.6 µg/ml Tm. (**D**) cER inheritance in SFA1 and SFA4 cells upon treatment with Tm, an ER stress inducer. SFA1 or SFA4 strains expressing Pho88-GFP were treated with control DMSO or Tm for 90 minutes and then examined by fluorescence microscopy. Representative images of GFP or overlay of DIC with GFP of class I, II, and III cells are shown. Yellow arrowheads indicate normal cER in the bud, and white arrows point to the discontinuous cER under the cortex of the daughter cell. Scale bars, 5 µm. (**E**) Quantification of the cER inheritance of the cell shown in (D). For each class, over 100 cells were analyzed. Results represent analyses from three independent experiments. Error bars denote standard deviation (SD). A one-tailed *t-test* was used to determine the significance of the difference between SFA1 and SFA4 cells with and without Tm treatment, p < 0.01, ** ; p < 0.05, * ; ns, not significant.

Remarkably, SFA4 cells displayed elevated GFP fluorescence even without Tm treatment, reaching levels comparable to those observed in Tm-treated WT or SFA1 cells. This strong basal activation indicates that increased ER membrane lipid saturation in SFA4 is sufficient to induce UPR activation independently of external ER stressors. Upon Tm treatment, GFP levels in SFA4 cells increased further, demonstrating that the UPR machinery remains responsive to additional ER stress (**Fig. 4A-B**). These results support the idea that ER lipid saturation perturbs ER homeostasis and activates the UPR independently of unfolded protein accumulation, likely through direct sensing of membrane lipid composition by UPR transducers Ire1 in yeast. To corroborate the findings from the UPRE-GFP reporter assay, we evaluated the growth phenotypes of WT and SFA strains under ER stress conditions (**Fig. 4C**). Under unstressed conditions, SFA1, SFA2, and SFA3 strains exhibited growth comparable to WT. In contrast, SFA4 cells showed slower growth, consistent with a basal stress state. Upon ER stress induction with tunicamycin (Tm), all strains except SFA4 displayed a growth defect. Interestingly, SFA4 cells exhibited improved growth in the presence of Tm relative to their untreated condition, and even outperformed other SFA strains under ER stress. These results suggest that Tm-induced proteotoxic stress activates the UPR in a way that may partially compensate for the lipotoxic defects in SFA4 cells.

To further investigate the unexpected behavior of SFA4, we investigated if/how ER stress alters the cortical ER (cER) inheritance when compared to that observed in unstressed SFA4 cells. In all classes of cells for SFA4, induction of ER stress with Tm restored cER inheritance, enabling all cells to properly inherit cER despite significant alterations in overall ER morphology (**Fig. 4D–E, *S2B***). Notably, unstressed SFA4 class II cells exhibited a pronounced decrease in cER inheritance (**Fig. 4E**, class II, gray bar). However, upon induction of ER stress with Tm, the reduced cER inheritance in SFA4 class II cells was restored to nearly normal levels (**Fig. 4D–E, *S2B***). This was in contrast to SFA1 cells; when we assessed cER inheritance across all three classes with ER stress, cER inheritance was blocked in class I, II, and III buds in a manner similar to that observed in wild-type cells (**Fig. 4D–E**). These findings suggest that elevated lipid saturation in SFA4 impacts ER inheritance efficiency, while ER stress-induced changes in lipid composition could facilitate improved cER inheritance. Alternatively, ER stress may alter the biophysical characteristics of yeast ER lipids in a way that enhances the effectiveness of cER inheritance.

### Impact of Inositol Depletion on Cortical ER Inheritance and ERAD Substrate Degradation

Outcomes of above studies have motivated us to test other types of lipid changes on ER inheritance. Inositol is a vital precursor for the synthesis of phosphatidylinositol (PI) in yeast. Previous research has demonstrated that inositol depletion activates the unfolded protein response (UPR) and leads to significant alterations in cellular phospholipid composition^38,39^. To investigate whether membrane alterations caused by inositol depletion affect ER quality control, we employed LipSat sensors (**Fig. *S1B-C***) to assess ER or INM membrane lipid saturation under inositol-depleted conditions (**Fig. 5A-B and *S3A-B***). Our analysis revealed that inositol deprivation does not cause major changes to lipid saturation levels at the ER or INM, suggesting that the observed effects on the ER homeostasis are not due to altered membrane saturation.

**Figure 5.**
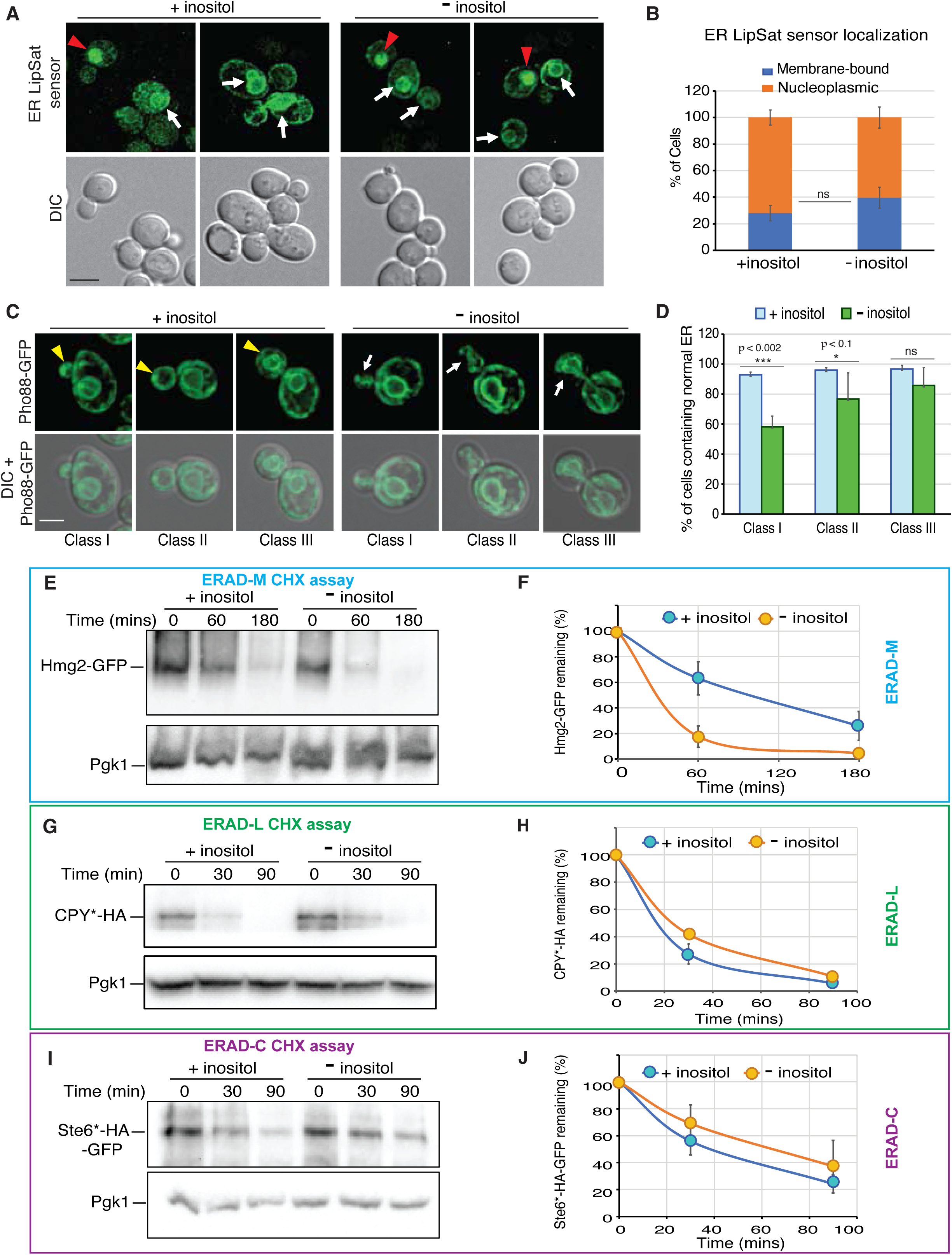
Inositol depletion affects cER inheritance and ERAD substrate degradation. (**A**) ER membrane lipid saturation levels in response to inositol depletion. Live cell imaging of wild-type cells expressing the plasmid-based ER LipSat sensor under conditions with or without 100 µM inositol supplementation. Red arrowheads indicate nucleoplasmic localization, and white arrows point to nuclear envelope localization. Scale bar, 5 µm. (**B**) Localization of the ER LipSat sensor was quantitated to assess lipid saturation levels. Sensor distribution was classified into two phenotypes: membrane-bound (indicative of unsaturated lipid) or nucleoplasmic (indicative of saturated lipids). Data are presented as mean values and standard deviations. A one-tailed *t-test* shows no significant difference for ER lipid saturation between the two growth conditions. (**C**) Inositol depletion leads to defects in cER inheritance. Wild-type cells expressing the ER membrane reporter, Pho88-GFP, were analyzed by fluorescence microscopy under normal or inositol-depleted conditions. Representative images show abnormal cER inheritance, with yellow arrowheads indicating normal cER formation in the bud and white arrows indicating defective cER. Scale bar, 5 µm. (**D**) Quantification of cER inheritance defects. More than 100 cells were analyzed under conditions of inositol supplementation (+Inositol) or depletion (-Inositol). Data represent the average of three independent experiments, with error bars indicating standard deviation (SD). Statistical significance of cER inheritance differences in wild-type cells grown with or without inositol was assessed using a one-tailed *t- test*. p < 0.002, *** ; p < 0.1, * ; ns, not significant. (**E**) Cycloheximide (CHX) chase degradation assay was performed for ERAD-M substrate, Hmg2-GFP, and analyzed by quantitating remaining Hmg2-GFP at each time point following the release from cycloheximide chase. (**F**) Quantification of ERAD-M (Hmg2-GFP) levels shows the percent of Hmg2-GFP remaining at each time point from the cycloheximide chase assay in panel (E). (**G**)-(**H**) Cycloheximide chase degradation assay was performed for ERAD-L substrate, CPY*-3xHA and analyzed by quantitating remaining CPY*-3xHA at each time point following the release from cycloheximide chase. (**I**)-(**J**) Cycloheximide chase degradation assay for ERAD-C substrate, Ste6*-HA-GFP, In all cases (E-H), Pkg1 was used for loading controls for all the quantitation. In all cases, means of percent remaining for each ERAD substrate from three biological replicates are plotted, with error bars representing standard deviation (SD).

We next examined the impact of inositol depletion on cER inheritance and ERAD pathways. Under inositol-depleted conditions, early-stage (Class I) cells successfully formed ER tubules emerging from the pnER, extending into the bud and anchoring at the bud tip. However, in later-stage (Class II) cells, while ER tubules persisted, cER formation was partially inhibited or delayed. By the final stage (Class III), despite proper pnER was established in the bud, cER failed to uniformly distribute along the bud cortex (**Fig. 5C, *S3C***). Quantitative analysis confirmed significant defects in cER inheritance due to inositol depletion, although these defects were less severe than those induced by tunicamycin (Tm)–mediated ER stress (**Fig. 5D**, in comparison to **Fig. 1D**).

To assess the influence of inositol depletion on ERAD, we measured the turnover of three distinct ERAD substrates. Under inositol-depleted conditions, degradation of the ERAD-M substrate Hmg2-GFP accelerated (**Fig. 5E-F**). In contrast, both the ERAD-L substrate CPY*-HA (**Fig. 5G-H**) and the ERAD-C substrate Ste6*-HA-GFP (**Fig. 5I-J**) exhibited mild stabilization, with only slight decrease in their degradation rates, suggesting a modest impairment in the breakdown of these substrates. Previous studies have reported that inositol depletion alters ER membrane phospholipid composition—specifically decreasing PI levels while elevating cardiolipin (CL) and phosphatidylglycerol (PG) levels^40–42^ (**Fig. *S4A***). These changes modulate overall membrane lipid homeostasis, which collectively enhances the degradation of ERAD-M substrates. In contrast, the degradation of ERAD-L and ERAD-C substrates remains largely unaffected, indicating that lipid perturbation under inositol depletion selectively impacts the ERAD-M pathway. To determine whether inositol depletion–induced membrane aberrations act synergistically with increased ER membrane lipid saturation, we assessed the growth of wild-type and SFA1–SFA4 strains on medium with or without inositol. Wild-type, SFA1, and SFA2 cells displayed no growth defects under inositol deprivation. However, the SFA3 strain, which grew a little slowly under normal conditions, showed pronounced sensitivity to inositol depletion, with severe growth inhibition at high temperature 37^0^C on inositol-depleted plates (**Fig. *S4B***, blue box). This trend became even worse with SFA4 at 30°C, -inositol, cell growth was diminished significantly (**Fig. *S4B***, red box). These results indicate a synthetic interaction between inositol depletion and elevated ER membrane lipid saturation.

### Deletion of specific phospholipid biosynthesis genes disrupts cER inheritance

In addition to studying the effects of inositol depletion, we investigated how disruption of key steps in phospholipid biosynthesis pathways affects endoplasmic reticulum (ER) morphology, cortical ER (cER) inheritance, and ER quality control. Specifically, we focused on genes involved in the synthesis of major ER phospholipids, including phosphatidylcholine (PC), phosphatidylethanolamine (PE), and diacylglycerol (DAG) and related intermediates, including *INO4, CHO2, OPI3, PAH1,* and *SPO7* (**Fig. 6A**). These genes represent key regulatory nodes in the CDP-DAG and Kennedy pathways, with prior lipidomic profiling available for their respective knockout strains^43–45^, providing valuable context to interpret the functional consequences of lipid imbalance on ER homeostasis.

**Figure 6:**
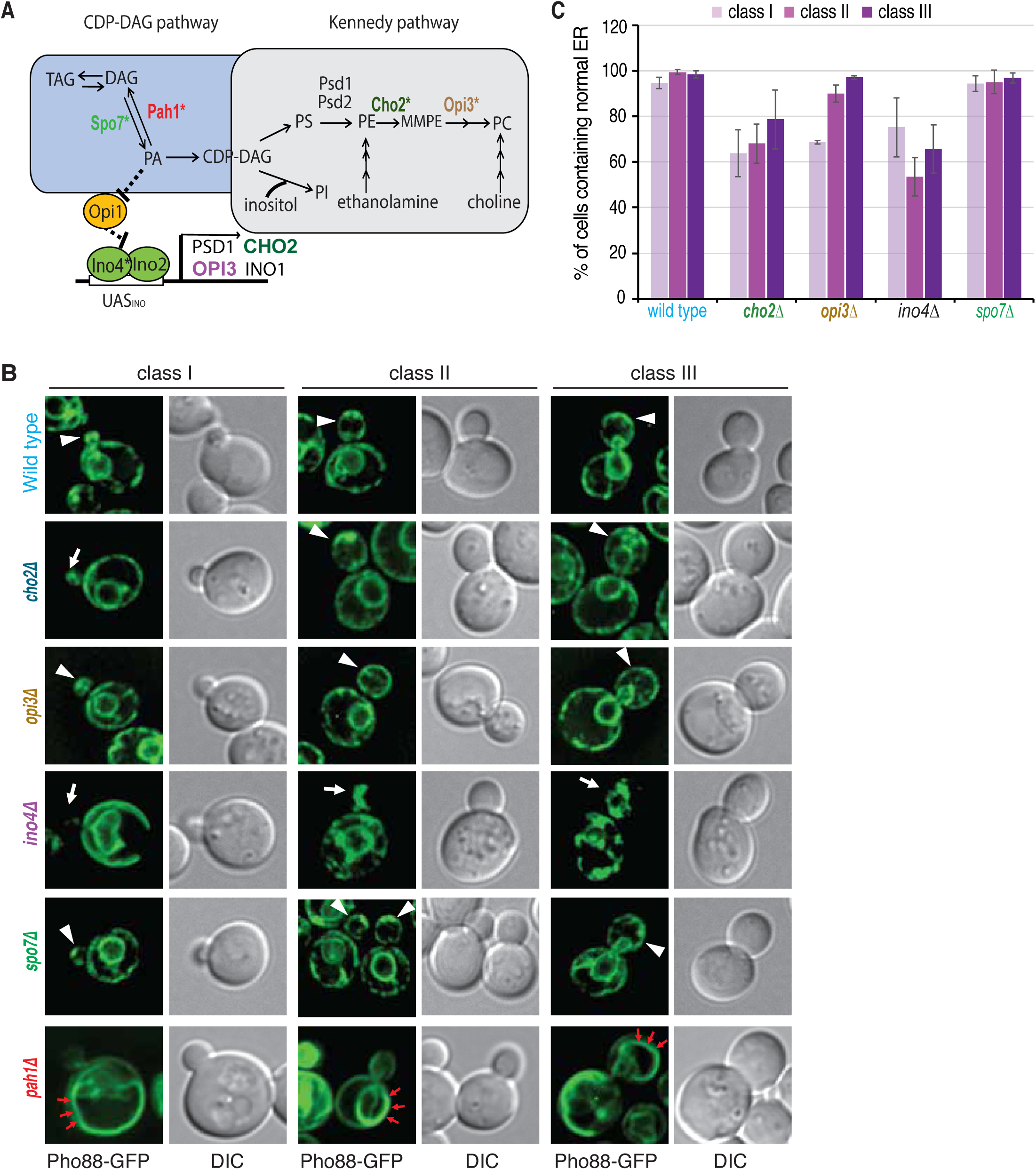
Impacts of Phospholipid synthesis deficient cells on ER structure and functions. (**A**) Abbreviated schematics of phospholipid biosynthesis pathways, specifically, the CDP-DAG pathway and the Kennedy pathway, were shown with some of the major intermediates include phosphatidic acid (PA), CDP-diacylglycerol (CDP-DAG), triacylglycerol (TAG), phosphatidylserine (PS), phosphatidylethanolamine (PE), phosphatidylcholine (PC), phosphatidylinositol (PI) as well as monomethyl phosphatidylethanolamine (MMPE). Key enzymes involved in these pathways include PSD1/PSD2 (PS to PE), CHO2/OPI3 (PE to PC via MMPE), and INO1/INO2/INO4 for inositol biosynthesis and regulation. The Kennedy pathway utilizes inositol to generate PI. PAH1, a phosphatidate phosphatase, converts PA to diacylglycerol (DAG), promoting TAG synthesis. SPO7, a regulatory subunit of the Nem1-Spo7 complex, activates Pah1 to control PA levels. Genes marked with an asterisk indicate those analyzed in their null mutant form (e.g., *opi3Δ*, *cho2Δ*, *spo7Δ*, *pah1Δ, ino4Δ*) to dissect pathway function. (**B**) Disruptions in specific lipid synthesis pathways result in cER inheritance defects or abnormal ER structures. Wild-type (WT), *pah1Δ*, *cho2Δ*, *opi3Δ*, *ino4Δ*, and *spo7Δ* cells expressing the ER membrane marker Pho88-GFP were examined by fluorescence microscopy. Three categories of cells are depicted for each strain based on the bud index. Arrowheads indicate normal cER inheritance in the bud, while arrows highlight abnormal cER inheritance. Red arrows mark aberrant ER structures. Scale bar, 5 µm. (**C**) Quantification of cER inheritance in the indicated strains. The percentage of cells containing cER in the daughter cell was determined (n > 100 per strain). Severe deformation of the ER structure in *pahΔ* cells prevented quantitation of cER inheritance.

As illustrated in **Fig. 6A**, *INO4* encodes a transcription factor that heterodimerizes with *INO2* to activate genes involved in multiple phospholipid biosynthesis steps, including *CHO1*, *CHO2*, *OPI3*, *INO1*, *CDS1*, and *PIS1*^46, 47^. In *ino4Δ* cells, we observed pronounced ER morphological defects, affecting both the cER and pnER throughout the cell cycle, resulting in a strong block in cER inheritance (**Fig. 6B-C, and *S5***).

In the Kennedy pathway, Cho2 and Opi3 act sequentially to methylate phosphatidylethanolamine (PE) to generate phosphatidylcholine (PC), two of the predominant phospholipids in ER membranes^48^. Individual deletion of *CHO2* or *OPI3* reduces PC synthesis^49, 50^, leading to mild but consistent defects in cER inheritance (**Fig. 6B-C, Fig. *S5***). PC and PE differ in their biophysical properties: PC’s cylindrical shape promotes stable lamellar bilayer formation and membrane fluidity, while PE’s cone shape favors negative curvature and membrane bending. Imbalances in the PC:PE ratio likely alter membrane organization, fluidity, and curvature, causing lipid bilayer stress that can activate ER stress pathways. Notably, these lipid perturbations may trigger the ER stress surveillance (ERSU) mechanism, which monitors ER integrity and delays ER inheritance to daughter cells under stress conditions. The activation of ERSU in *opi3Δ* and *cho2Δ* strains likely contributes to their observed cER inheritance defects.

The lipid phosphatase Pah1 and its activator Spo7 regulate conversion of phosphatidic acid (PA) to DAG, a metabolic branch-point essential for both phospholipid and triacylglycerol (TAG) synthesis. In *spo7Δ* mutants, phospholipid levels are elevated while TAG levels decline, yet cER inheritance remains normal (**Fig. 6B-C, Fig. *S5***). Conversely, *pah1Δ* mutants exhibited pronounced ER structural abnormalities. In wild-type cells, Pho88-GFP localized to the typical cER and pnER surrounding well-defined nuclei, as confirmed by DAPI staining (data not shown). In contrast, *pah1Δ* cells displayed disrupted Pho88-GFP distribution characterized by a fragmented and disorganized ER network (**Fig. 6B, Fig. *S5***). DAPI staining in *pah1Δ* cells revealed irregular perinuclear ER morphology that correlated with the disturbed ER structure (data not shown). Differential interference contrast (DIC) images confirmed normal overall cell morphology in both strains. These observations are consistent with previous reports of ER sheet proliferation and expansion in *pah1Δ* mutants^51^, and thus, overall morphology of the ER in both mother and daughter cells were significantly different which hindered clear quantitative analysis of cortical ER inheritance. These findings suggest that loss of PA phosphatase activity disrupts ER morphology through mechanisms extending beyond changes in phospholipid to triacylglycerol ratios. Together, these results emphasize the critical role of phospholipid biosynthesis and homeostasis in maintaining ER morphology, facilitating cER inheritance, and preserving ER functional integrity.

### Impact of lipid imbalance on ER-associated degradation (ERAD)

To assess the effects of lipid perturbations on protein quality control, we examined degradation of substrates from the three branches of ER-associated degradation (ERAD). In ***opi3Δ*** cells, Hmg2-GFP (ERAD-M substrate) was strongly stabilized, indicating impaired degradation of membrane-embedded proteins (**Fig. 7A–B, *S6A–B***). This suggests that the altered membrane lipid composition in *opi3Δ* cells may disrupt key steps such as substrate recognition, retrotranslocation, or dislocation, all of which depend on the proper membrane environment. By contrast, *cho2Δ* cells displayed degradation kinetics of Hmg2-GFP that were comparable to wild-type, suggesting that partial PC depletion does not substantially interfere with ERAD-M, or that compensatory mechanisms may be at play (**Fig. *S6A-B***). In the case of the luminal substrate CPY*-HA, both *opi3Δ* and *cho2Δ* cells showed clear stabilization (**Fig. 7C-D, *S6C-D***), indicating that ERAD-L (the branch handling misfolded proteins in the ER lumen) is sensitive to disruptions in PC levels. These defects could arise from lipid-induced perturbations in the folding environment, luminal chaperone activity, or dislocation machinery. In contrast, the degradation of Ste6*-HA-GFP was unaffected in both mutants (**Fig. 7E-F**, ***S6E-F***), indicating that ERAD-C, which targets cytoplasmic misfolded domains, remains intact. This suggests that the cytosolic arm of the quality control machinery is either independent of ER lipid composition or more robust against moderate membrane perturbations.

**Figure 7:**
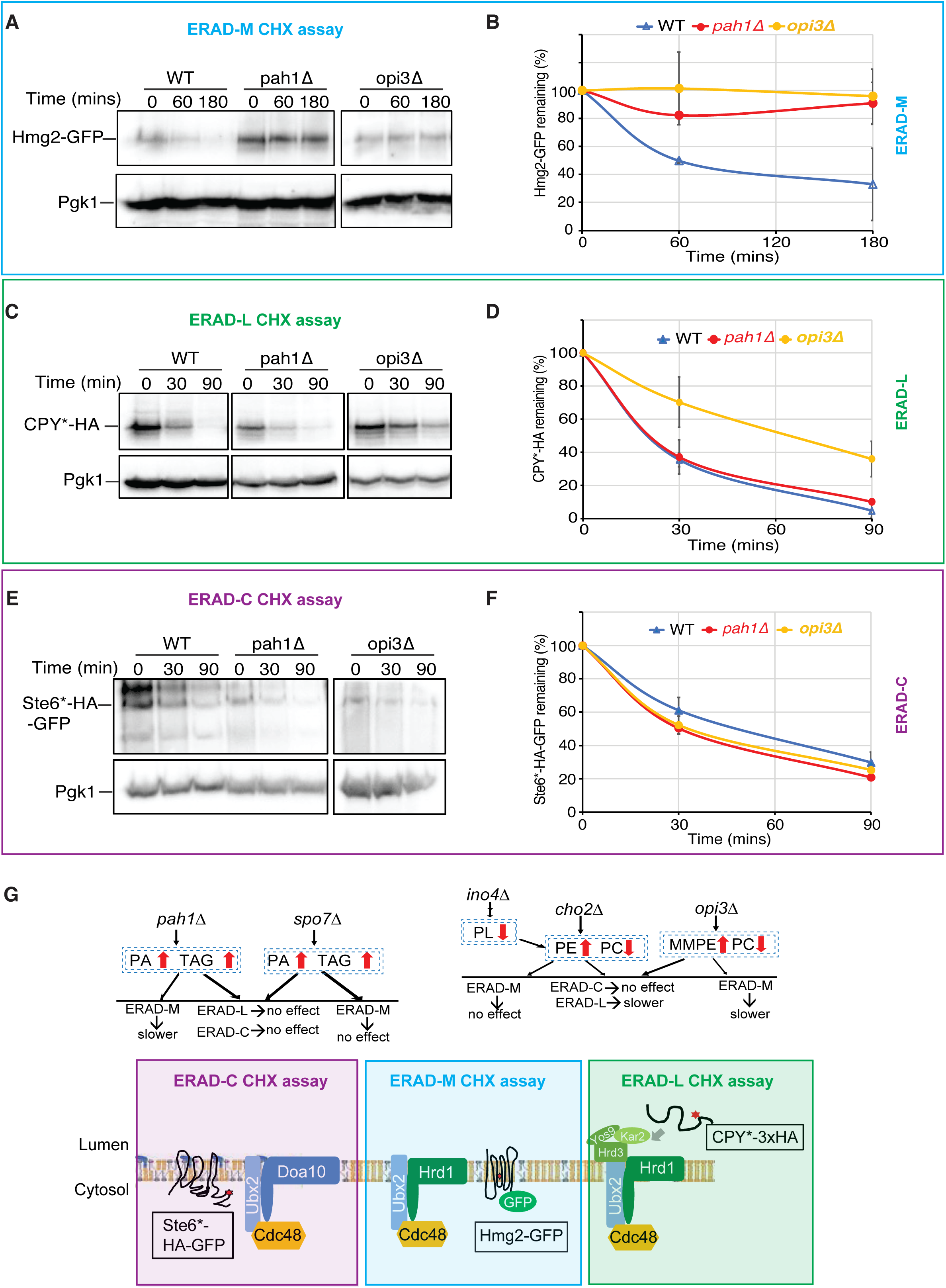
Effects of lipid synthesis pathway mutants on ERAD substrate degradation. (**A**)(**C**)(**E**): Cycloheximide chase degradation assay for the ERAD substrate was performed in wild-type (WT), *pah1Δ* and *opi3Δ* cells with chromosomally integrated (A) the ERAD-M substrate, Hmg2-GFP; (C) the ERAD-L substrate, CPY*-3xHA; (E) the ERAD-C substrate, Ste6*-HA-GFP; (**B**)(**D**)(**F**): Quantification of (A),(C) and (E), respectively. For all assays, total protein levels were assessed using anti-Pgk1 blots as a loading control. Representative immunoblots from at least three independent biological replicates are shown. The means of each ERAD substrate remaining were plotted, with error bars representing the standard deviation (SD). (**G**) Impact of lipid biosynthesis gene deletions on ERAD substrate degradation in yeast. This schematic summarizes the data from Fig. 7 and Fig. S6 how deletions of lipid metabolism genes—*cho2Δ*, *opi3Δ*, *spo7Δ*, *pah1Δ*, and *ino4Δ*—affect the degradation of substrates from the three branches of ER-associated degradation (ERAD): ERAD-C (Ste6*-HA-GFP), ERAD-M (Hmg2-GFP), and ERAD-L (CPY*-3xHA).

To further dissect the relationship between lipid homeostasis and ERAD, we examined *ino4Δ* cells, which have broader disruptions in phospholipid biosynthesis due to loss of a key transcriptional regulator. These cells behaved similarly to *cho2Δ*, showing no degradation defects for Hmg2-GFP or Ste6*-HA-GFP, but impaired degradation of CPY*-HA (**Fig. S6A-F**). This result underscores the particular sensitivity of ERAD-L to changes in overall phospholipid biosynthesis and supports the notion that membrane integrity is critical for luminal protein quality control. Next, we examined *pah1Δ* and *spo7Δ* mutants, which exhibit elevated phospholipid biosynthesis and reduced triacylglycerol (TAG) accumulation. Interestingly, both mutants maintained efficient degradation of CPY*-HA and Ste6*-HA-GFP (**Fig. 7C-F**, ***S6C-F***), indicating that ERAD-L and ERAD-C remain largely functional. However, in *pah1Δ* cells, Hmg2-GFP was significantly stabilized (**Fig. 7A-B**, ***S6A-B***), suggesting a selective impairment of ERAD-M. This observation is consistent with previous reports of dramatic ER morphological changes in *pah1Δ*, including ER sheet expansion, which may disrupt membrane protein surveillance by affecting membrane fluidity, curvature, or the spatial organization of ERAD machinery.

Together, these findings demonstrate that distinct perturbations in lipid metabolism differentially affect ER-associated degradation pathways depending on the subcellular localization of the misfolded domain. Specifically, membrane protein degradation is highly sensitive to membrane structure alterations (*opi3Δ*, *pah1Δ*), luminal protein degradation is particularly affected by reduced PC or global phospholipid synthesis (*opi3Δ*, *cho2Δ*, *ino4Δ*), while degradation of cytosolic misfolded proteins is largely preserved. These data highlight the critical role of phospholipid homeostasis in maintaining ER quality control functions and suggest that ERAD branches are differentially tuned to the biophysical and biochemical properties of the ER membrane.

## Discussion

The endoplasmic reticulum (ER) is a highly dynamic organelle central to protein folding, lipid synthesis, and the faithful inheritance of organelles during cell division. Our findings demonstrate that ER lipid homeostasis—encompassing saturation level, acyl chain composition, and phospholipid headgroup identity—is fundamental for maintaining ER architecture, dynamics, and proteostasis. Perturbations to lipid balance, whether arising from genetic disruption of biosynthetic pathways or environmental challenges such as inositol depletion, compromise ER integrity in distinct, branch-specific ways across ER-associated degradation (ERAD) pathways.

Lipid saturation emerged as a central determinant of ER membrane biophysical properties and function. Increased saturation, as modeled in SFA4 cells, reduced membrane fluidity and curvature, impairing cortical ER (cER) inheritance and producing structural defects during polarized growth. These observations support models in which dynamic membrane remodeling is essential for proper ER partitioning during budding. In contrast, strains with shorter or more unsaturated acyl chains, such as SFA1, showed much milder phenotypes, suggesting that acyl chain flexibility can buffer against lipid-induced stress and preserve ER dynamics.

Altered membrane composition also shaped the efficiency of ER quality control. Saturated membranes enhanced degradation of certain membrane and cytosolic ERAD substrates, possibly by making misfolded proteins less stable within a more rigid tightly packed lipid bilayer, thereby affecting recognition and processing^52^. This supports models in which the biophysical properties of the lipid bilayer modulate substrate conformations and their accessibility to degradation machinery. Yet, ERAD-L substrates were spared, highlighting differences in the sensitivity of ERAD branches to changes in membrane composition. Indeed, the divergent properties of the SFA strains offer mechanistic insight. SFA1 cells maintain a largely wild-type phospholipid profile with shorter chains, whereas SFA4 cells accumulate fully saturated phospholipids that generate rigid membranes^36^. In wild-type cells such saturated lipids are present only as part of a more diverse lipid pool, buffering their impact. As summarized in Fig 3G, these compositional differences are likely substrate-specific in their effects. For membrane-associated substrates such as Hmg2- GFP (ERAD-M) and Ste6*-HA-GFP (ERAD-C), increased saturation may impede recognition, ubiquitination, or retrotranslocation^8, 53^. Each of these steps depends on dynamic interactions between misfolded domains and membrane-embedded quality control factors, processes that are highly sensitive to bilayer rigidity. In contrast, CPY*-HA, a soluble luminal substrate requiring recognition by chaperones such as BiP, bypasses reliance on the membrane environment and is therefore less directly affected by lipid saturation, although ERAD-L substrates require to pass through the lipid bilayer to reach to the proteosomes present in cytoplasm.

The differential effects of *cho2Δ* and *opi3Δ* on ERAD-M and ERAD-L highlight the importance of lipid composition in regulating ER proteostasis. Although both mutants reduce phosphatidylcholine (PC) synthesis, only *opi3Δ* significantly impairs degradation of ERAD-M substrates. This suggests that the accumulation of MMPE—a cone-shaped lipid intermediate specific to the *opi3Δ* block—may disrupt ER membrane architecture by introducing excess curvature stress, thereby destabilizing membrane-resident degradation pathways. In contrast, *cho2Δ* cells, which do not accumulate MMPE, preserve ERAD-M function despite reduced PC levels, indicating that the quality and shape of accumulated lipids, rather than overall PC abundance, dictate ERAD-M efficiency. Notably, both mutants impair ERAD-L substrate degradation, reflecting the greater sensitivity of luminal quality control pathways to disruptions in lipid homeostasis.

Broader metabolic disruptions reinforced these trends. In ***ino4Δ*** cells, loss of a transcriptional regulator with broad control over phospholipid synthesis selectively impaired ERAD-L, while preserving ERAD-M and ERAD-C. Similarly, ***pah1Δ*** cells, which accumulate phospholipids and exhibit dramatic ER expansion, displayed selective defects in ERAD-M—likely reflecting how large-scale morphological disorganization disrupts the spatial coordination required for membrane protein surveillance.

Together, these findings reveal that ER lipid homeostasis exerts a profound influence on both organelle architecture and protein quality control. Lipid saturation and headgroup balance dictate membrane flexibility, curvature, and spatial organization, which in turn determine how efficiently different ERAD branches recognize and process misfolded clients. The results underscore that proteostasis cannot be viewed independently of lipid metabolism; instead, membrane composition acts as an active regulator of quality control pathways, tailoring ER architecture and function to the metabolic state of the cell.

### Experimental Procedures

#### Yeast and Plasmids Methods

Yeast cultures were grown at 30 °C in standard growth medium as previously described^54^. Plasmids were introduced into yeast using the lithium acetate transformation method^55^. A complete list of yeast strains and plasmids used in this study is provided in Tables S1 and S2, respectively. Gene deletions and epitope tagging were performed using PCR-based homologous recombination as described by Longtine et al.^56^. Plasmids pRH469, pRH1377, pRH2058, and pRH2997 were generously provided by the Randy Hampton lab. Plasmids pRH1377, pRH2058, or pRH2997 were directly transformed into the indicated yeast strains. Plasmid pRH469 was linearized with StuI and integrated into the URA3 locus of the specified strains. ER localization was monitored using Pho88-GFP, a GFP-tagged ER membrane protein^57^.

For inositol depletion experiments, yeast strains were initially grown on inositol-containing plates. Cells were then cultured overnight in inositol-free medium (prepared using yeast nitrogen base without amino acids and inositol) to reach mid-exponential phase. Cultures were freshly diluted into inositol-free medium and incubated for an additional 4–5 hours prior to fluorescence imaging or cycloheximide chase assays. In control experiments, 100 μM inositol was added to the inositol-free medium.

Fatty acid–supplemented growth media were prepared as previously described^37^. Briefly, standard SC medium was modified to contain 0.1% glucose and 1.5% Brij L23 (Sigma-Aldrich), and supplemented with 16 mM fatty acids (oleic acid or linoleic acid; Sigma-Aldrich). Control media were prepared identically but without the addition of fatty acids.

### Growth test

Yeast growth was assessed using a serial dilution spot assay. Yeast strains were cultured overnight in standard growth medium at 30°C with shaking until they reached mid-log phase. Cultures were then adjusted to an OD_600_ of 1.0, and a series of 5-fold serial dilutions were prepared in sterile water. Five microliters of each dilution were spotted onto appropriate solid agar plates using a multichannel pipette. Plates were incubated at the indicated temperatures and time points, and growth was documented by imaging. Representative images were taken after the appropriate incubation period to compare growth differences between strains.

### Fluorescence microscopy imaging

Fluorescence microscopy was performed using a Zeiss Axiovert 200M fluorescence microscope (Carl Zeiss MicroImaging) equipped with standard Zeiss filter sets for DAPI, FITC/GFP and TRITC/RFP. Yeast cells were grown to mid-log phase in the indicated media, harvested by centrifugation, and resuspended in phosphate-buffered saline (PBS) prior to imaging. Live-cell imaging was performed using a 100× 1.3 NA oil immersion objective, and images were captured with a monochrome digital camera (Axiocam; Carl Zeiss MicroImaging). For each field, z-stacks spanning 5 µm were collected at 0.2 µm step intervals. Following image acquisition, the stacks were deconvolved and projected to generate 2D images for subsequent quantitation. Exposure times and imaging settings were kept constant across all samples to ensure comparability. Image acquisition, deconvolution, and analysis were carried out using Zeiss Zen software.

To quantify cortical ER (cER) inheritance, more than 100 budded cells were analyzed per condition. Cells were classified into three categories based on bud index (the ratio of bud size to mother cell diameter), and the presence or absence of cER in the bud was scored, as previously described ^30, 58^.

Nuclear DNA was stained with DAPI to visualize nuclear position and facilitate identification of perinuclear ER localization in live cells. Cells were incubated with 1 µg/mL DAPI in PBS for 5–10 minutes at room temperature, followed by a PBS wash prior to imaging.

### IET tubule measurement protocols

To quantify initial ER tubules (IET)—a recently introduced term describing nascent ER tubules that emerge from the perinuclear ER and extend into the bud^59^—fluorescence microscopy images were analyzed using ImageJ software. Yeast cells expressing fluorescently tagged ER markers were grown to mid-log phase in appropriate media and imaged using a Zeiss fluorescence microscope equipped with a 100× oil immersion objective. Images were acquired under identical exposure settings to ensure consistency across samples.

For image analysis, raw fluorescence images were opened in ImageJ, and background subtraction was performed using the "Subtract Background" function. Tubules were manually traced using the “Freehand Line” tool or the “Segmentation” tool for more precise edge detection. The “Measure” function was then used to determine tubule length in pixels, which was converted to micrometers based on the microscope’s calibration settings. At least 50 tubules per condition were measured from multiple fields of view. To quantify tubule fluorescence intensity, tubules were selected as regions of interest (ROIs) using the “Freehand Selection” or “Polygon Selection” tool, ensuring that the entire tubule was encompassed.

The “Measure” function was then used to determine the mean fluorescence intensity and area of the ROI. Background fluorescence was subtracted by measuring an adjacent non-fluorescent region. At least 50 tubules per condition were analyzed across multiple fields of view. Statistical analysis was conducted using Microsoft Excel, with data analyzed using a student’s t-test. Results are presented as mean ± standard error of the mean (SEM).

### UPRE-GFP assay and quantitation

The unfolded protein response element (UPRE)-GFP reporter assay was used to assess UPR activation. Yeast cells carrying the UPRE-GFP reporter plasmid were grown to mid-log phase in selective media at 30°C. Cells were then either left untreated or subjected to ER stress by treatment with 1ug/ml tunicamycin for 90mins. Following treatment, cells were harvested by centrifugation, washed with phosphate-buffered saline (PBS), and resuspended in PBS for fluorescence microscopy imaging.

Fluorescence images were captured using a Zeiss fluorescence microscope with a 100× oil immersion objective. To quantify GFP intensity, images were analyzed using ImageJ software. Regions of interest (ROIs) were manually selected to measure fluorescence intensity within individual cells. Background fluorescence was subtracted, and mean fluorescence intensity (MFI) was calculated for each condition. Data were normalized to cell count and analyzed for statistical significance using Excel t-test.

### Cycloheximide Chase Degradation Assay

The degradation of epitope-tagged proteins was assessed using a cycloheximide chase assay, as described by Gardner et al. (1998)^60^. Briefly, yeast cultures grown to the logarithmic phase were treated with 100 μg/ml cycloheximide to inhibit protein synthesis. Samples were then collected at various time points post-treatment, lysed, and subjected to immunoblotting to evaluate protein degradation, following the protocol outlined by Gardner et al. (1998)^60^. GFP-tagged and HA-tagged proteins were detected using anti-GFP (Roche, catalogue number 11814460001) and anti-HA monoclonal antibodies (16B12, Covance, catalog number MMS-101P-500). Pgk1 is used as loading control and anti-Pgk1 is provided from Invitrogen (catalog number 459250).

## Acknowledgment

We thank the members of the Niwa, Kohler, Hampton, Budin, and Neal labs for their valuable discussions and insights. We are also grateful to Alwin Kohler, Itay Budin, Randy Hampton, Sonya Neal, and Joel Goodman for generously providing plasmids, yeast strains, and reagents. This work was supported by the Paul Allen Foundation Investigator Award and NIH (RO1GM087415), and CRCC to M.N.

---

**Figure S1:**
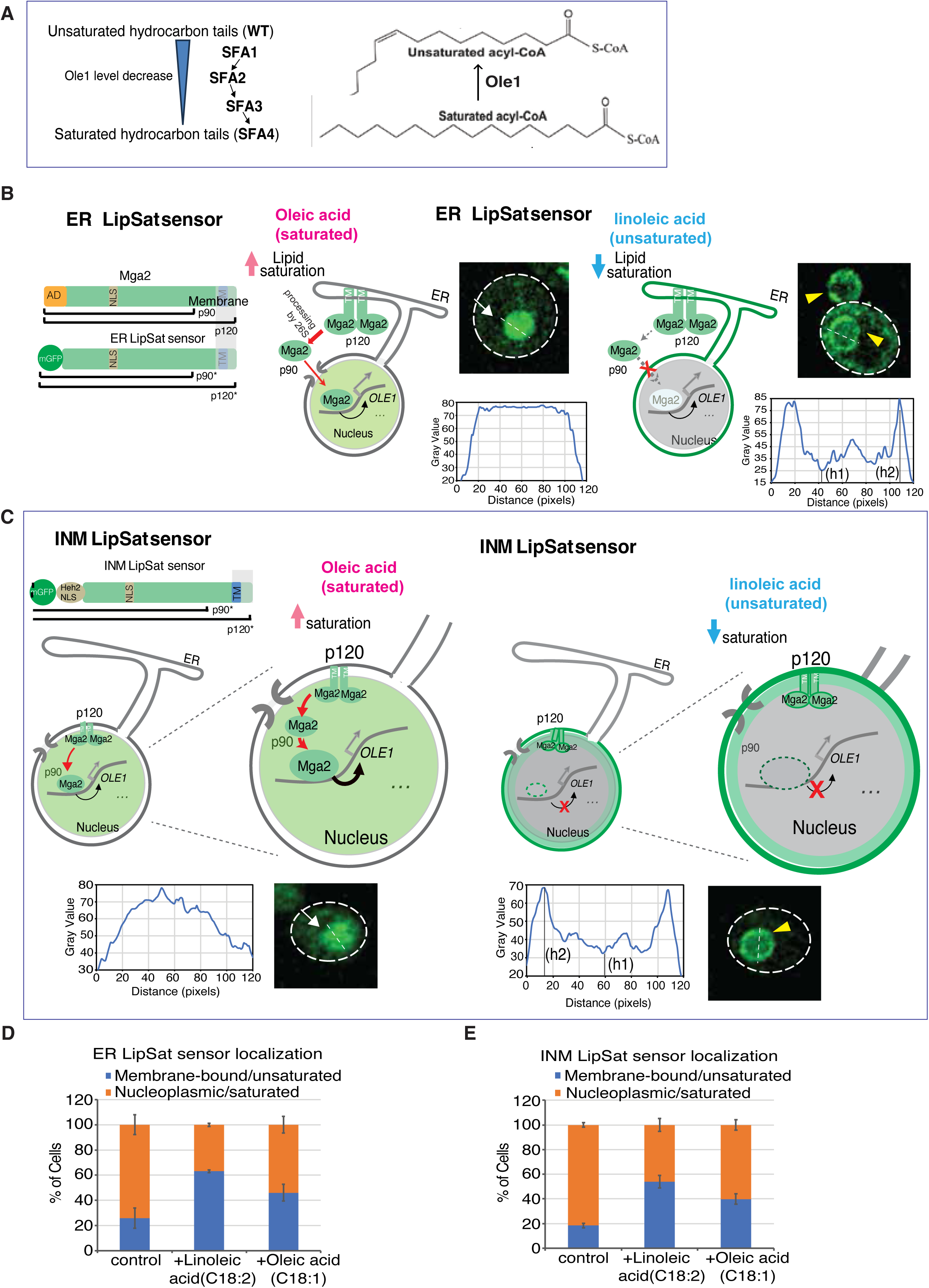
Extent of Lipid saturation is measured by ER LipSat and INM LipSat sensors. (**A**) Schematic illustrating the modulation of Ole1 expression levels in SFA1–SFA4 yeast strains^36^. OLE1 expression is placed under the control of different promoters to progressively reduce desaturase activity and elevate membrane lipid saturation. Drawing was modified from Venkatraman et al., 2023^36^. (**B**)(**C**) The ER LipSat (B) and INM LipSat (C) sensors^37^ enable assessment of fatty acid (FA) saturation in the ER membrane and the inner nuclear membrane (INM). (B) Mga2, an ER membrane localized transcription factor, undergoes proteolytic cleavage when membrane saturation increases, allowing its N-terminal portion to translocate into the nucleus to activate transcription. Therefore, the relative distribution of nuclear versus ER-localized Mga2 serves as a readout for ER membrane saturation. (**C**) Similarly, the INM LipSat sensor reports on saturation at the INM (or perinuclear ER). Mga2 localized to the INM by the Heh2-Nuclear localization signal (NLS) undergoes cleavage in response to elevated FA saturation, with its N-terminal GFP-tagged fragment released into the nucleoplasm. Diagram was adapted from Romanauska et al., 2021^37^. Without elevated FA saturation, INM LipSat stays on the INM. For each case, line plots showing GFP levels and localization of ER LipSat and INM LipSat sensors. GFP fluorescence was quantified along a line drawn across the center of the nucleus. The quantification method was adapted from Romanauska et al., 2021^37^. Fluorescence intensity profiles with a single central peak were classified as nucleoplasmic. For profiles with two or more prominent peaks, the lowest valley intensity (h1) and highest peak intensity (h2) were measured, and the ratio h1:h2 was calculated. A ratio > 0.5 was classified as nucleoplasmic, whereas a ratio < 0.5 indicated membrane associated localization. Wild-type cells (BY strain background) expressing plasmid-based ER LipSat sensor (MNY3371) or INM LipSat sensor (MNY3372) were grown in special medium with 0.1% glucose plus 1.5% Brij L23 solution supplemented with 16mM of the indicated fatty acid. Yellow arrowheads indicate membrane-bound sensor localization, and white arrows mark nucleoplasmic localization. Scale bar, 5 μm. (**D**) and (**E**) Quantification of ER LipSat and INM LipSat sensor localization from (B) and (C). Cells were categorized based on sensor localization as membrane-bound or nucleoplasmic. Over 100 cells per strain were analyzed.

**Figure S2:**
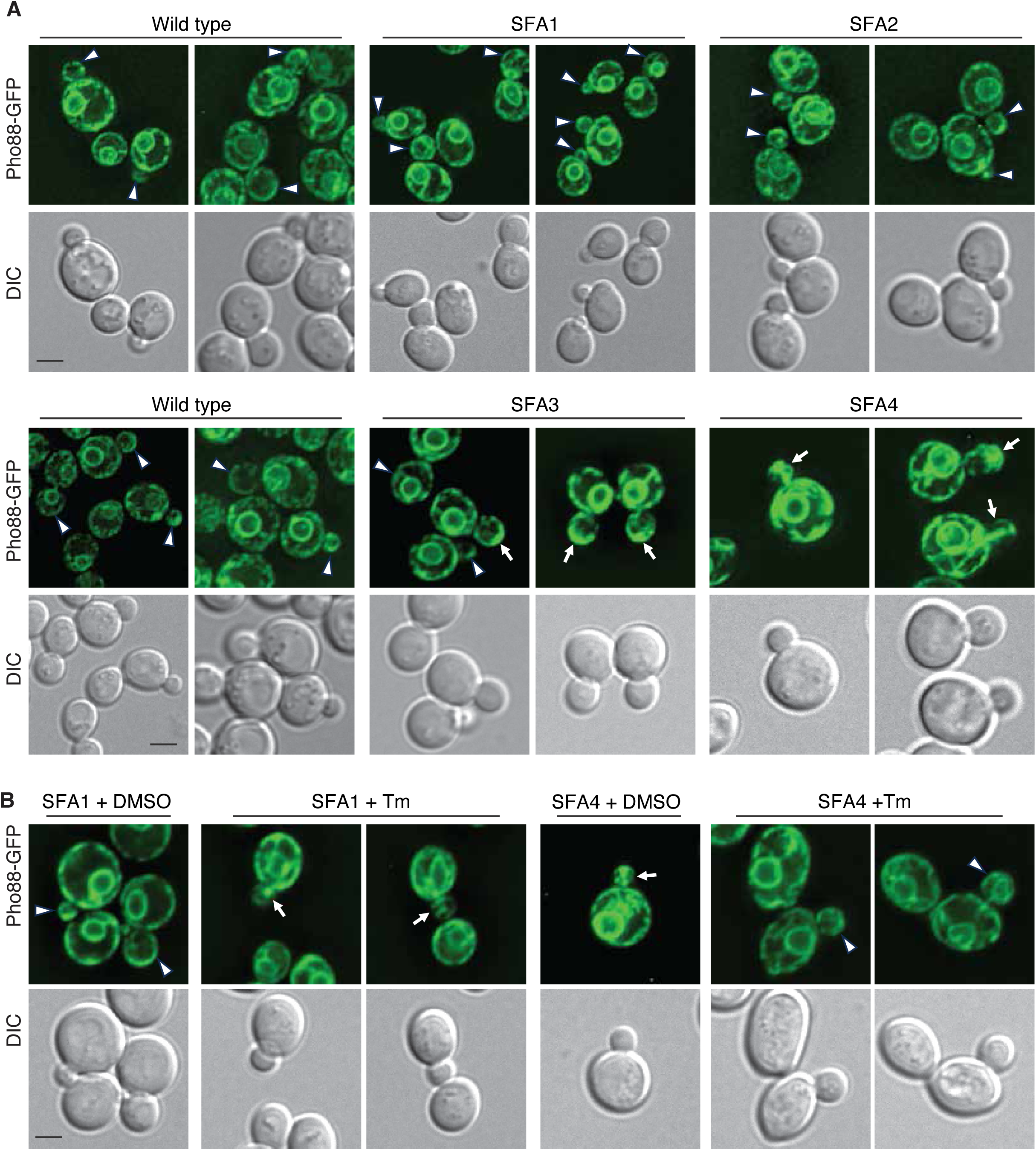
Effects of Lipid Saturation level on cER inheritance with or without ER stressor Tm treatment. (**A**) Fields of cells used for quantitation of cER inheritance in wild-type and SFA1–SFA4 cells shown in Figure 1D and 1E. Representative images of wild-type and SFA1–SFA4 cells expressing the ER membrane marker Pho88-GFP. Two different fields per specific strain are shown. Arrowheads indicate normal cER in daughter cells, while arrows mark examples of defective cER. Scale bar, 5 μm. (**B**) Tunicamycin (Tm) disrupts cER inheritance in SFA1 cells but rescues cER inheritance defects in SFA4 cells. SFA1 and SFA4 cells expressing Pho88-GFP were treated with 1 μg/ml Tm or DMSO (control) for 90 min before imaging. Two representative fields are shown. Arrowheads indicate examples of normal cER in daughter cells, while arrows mark defective cER. Scale bar, 5 μm.

**Figure S3:**
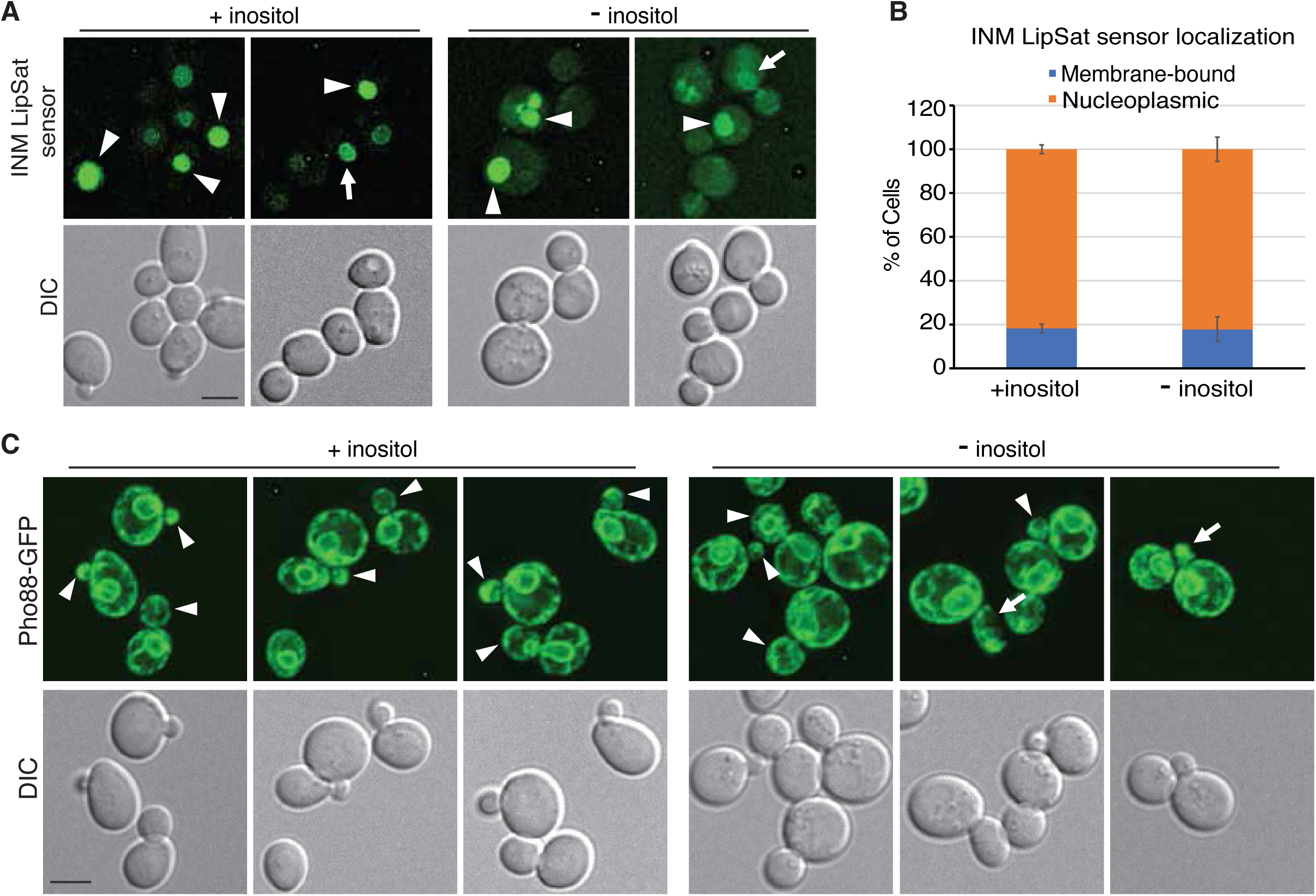
Effects of inositol depletion on the extent of FA saturation by localization of INM LipSat sensor and cER inheritance. (**A**) INM lipid saturation levels in response to inositol depletion. Live-cell imaging of wild-type cells expressing the plasmid-based INM LipSat sensor under conditions with or without 100 µM inositol supplementation. Arrowheads indicate FA saturation of the INM as INM LipSat sensor localized to nucleoplasmic localization, while arrows point to membrane-bound localization. Scale bar, 5 µm. (**B**) Quantification of INM LipSat sensor localization in panel (A). Sensor localization was classified as either membrane-bound or nucleoplasmic. Data represent mean values ± standard deviation (SD). (**C**) Inositol depletion leads to defects in cER inheritance. Wild-type cells expressing the ER membrane marker Pho88-GFP were examined by fluorescence microscopy under normal or inositol-depleted conditions. Representative fields are shown. Arrowheads indicate normal cER inheritance in the bud, while arrows highlight defective cER. Scale bar, 5 µm.

**Figure S4.**
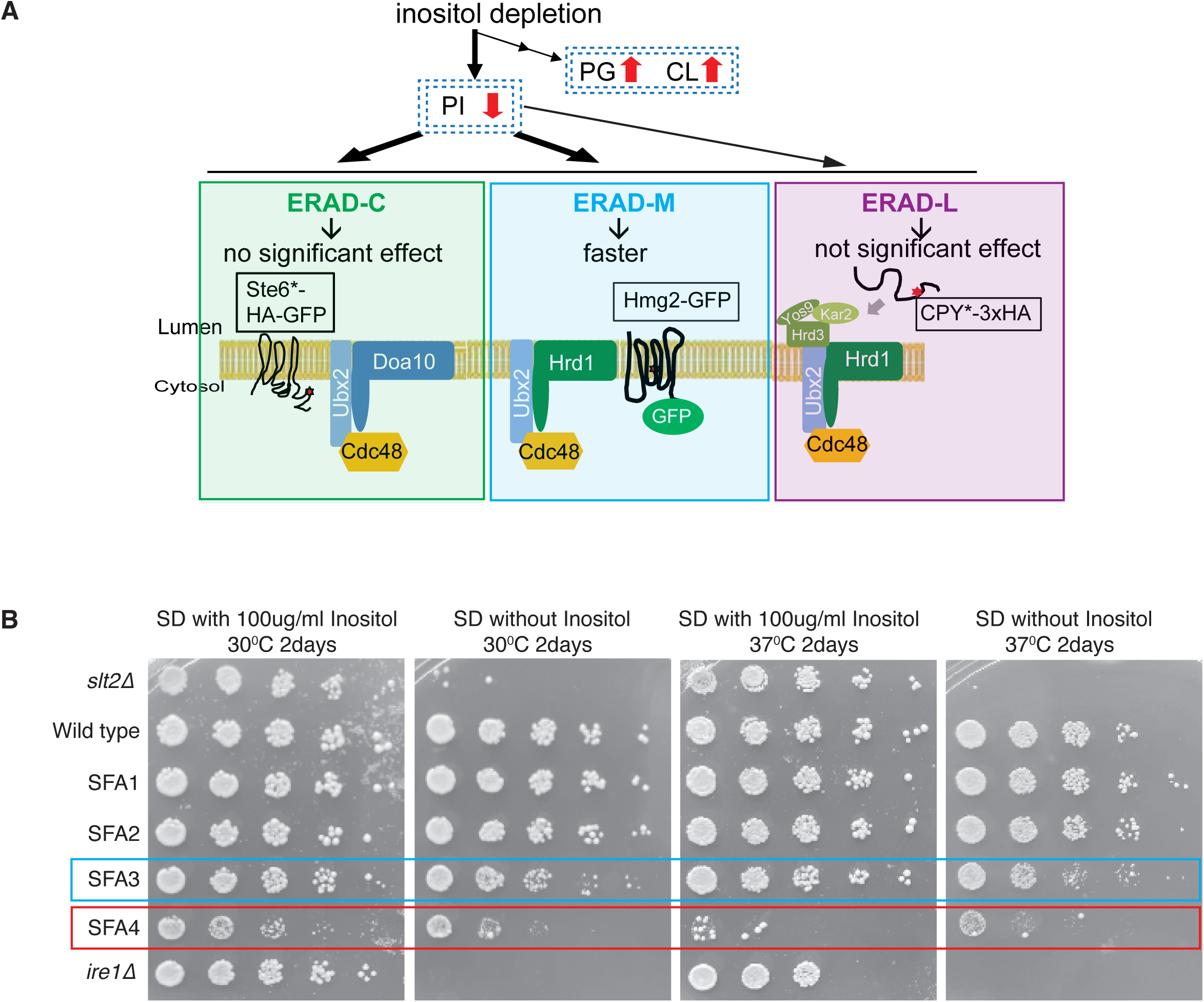
Synthetic growth phenotype of inositol depletion with membrane lipid saturation mutants SFA1 or SFA4 strains. (**A**) Cartoon summary illustrating the impact of inositol depletion on ER-associated degradation (ERAD) pathways. Under inositol-depleted conditions, degradation of ERAD-M substrates (e.g., Hmg2-GFP) is accelerated, while ERAD-C substrate (e.g., Ste6*-HA-GFP) degradation or ERAD-L substrate degradation (e.g., CPY*-3xHA) is not significantly affected. Substrate-specific E3 ubiquitin ligases and associated components are indicated: Hrd1 and Hrd3 for ERAD-L, Doa10 for ERAD-C, and Hrd1 for ERAD-M. Cdc48 and Ubx2 are involved in retrotranslocation and extraction for multiple ERAD branches. PI, PG and CL denote altered lipid environments resulting from inositol depletion, which may underlie differential effects of membrane alteration on substrate processing. (**B**) Growth phenotypes of SFA1 to SFA4 yeast in inositol-depleted conditions. Serial dilutions of overnight cultures from the indicated strains were plated on minimal synthetic plates with or without 100 µg/ml inositol and grown for 2 days at 30°C or 37°C. The blue and red boxes highlight the impaired growth of SFA3 and SFA4 strains under inositol depletion.

**Figure S5.**
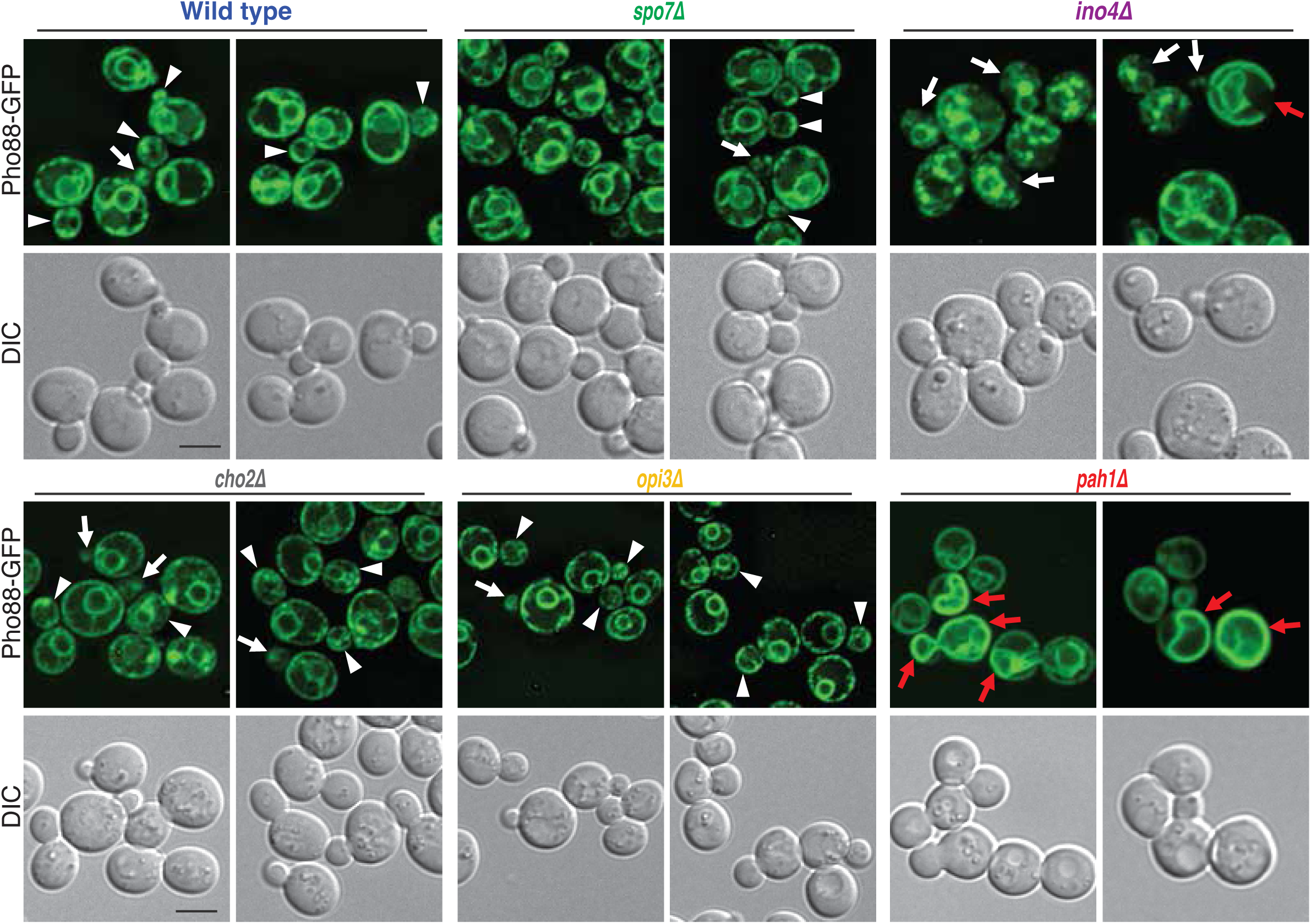
Supplementary data for Figure 6. Representative fields of wild-type (WT), *pah1Δ, cho2Δ, opi3Δ, ino4Δ,* and *spo7Δ* cells expressing Pho88-GFP are shown. Arrowheads indicate normal cER inheritance in the bud, arrows highlight defective cER inheritance, and red arrows mark aberrant ER structures. Scale bar, 5 µm.

**Figure S6:**
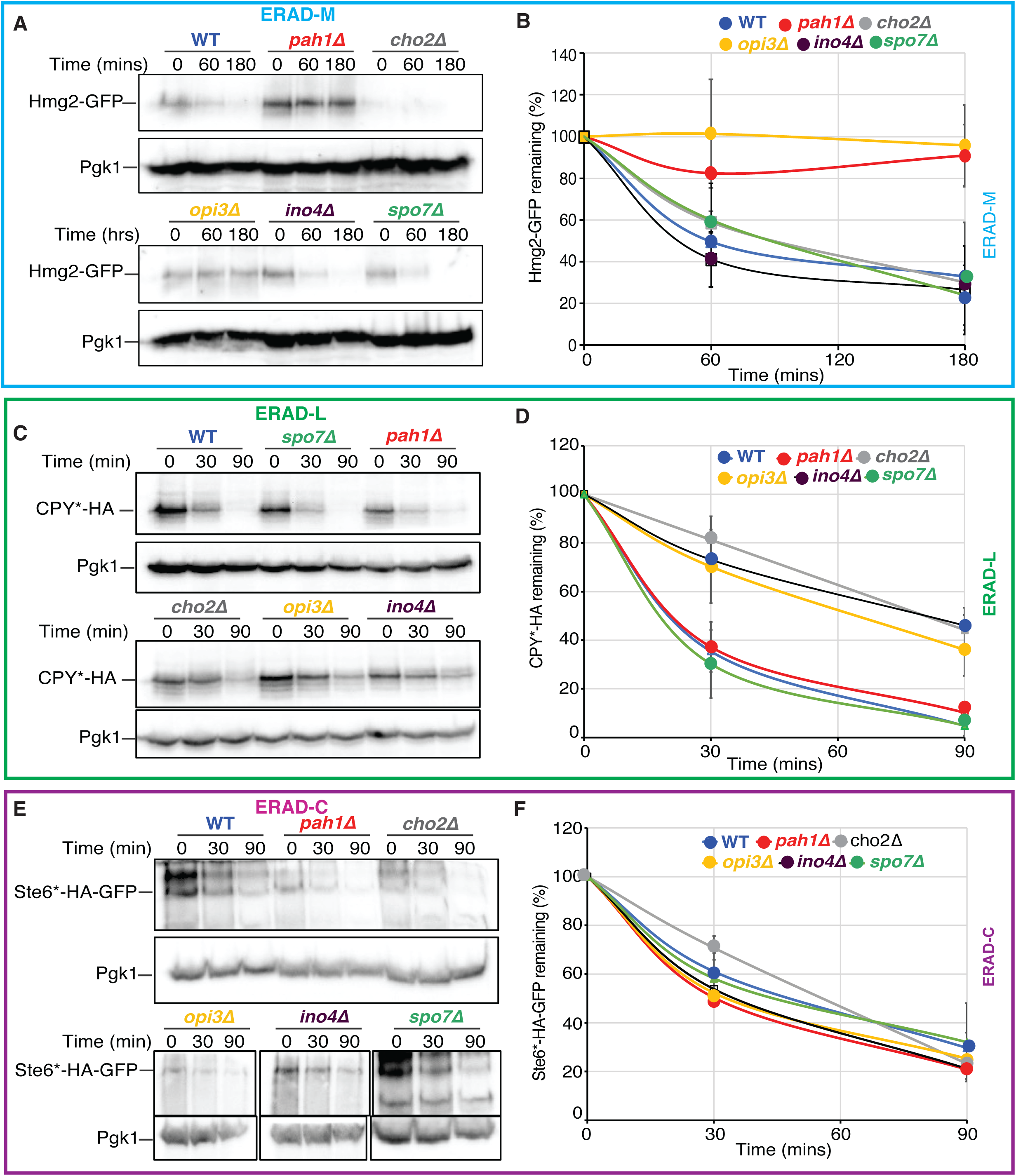
Effects of lipid synthesis pathway mutants on ERAD substrate degradation. (**A**) Cycloheximide chase degradation assay for the ERAD-M substrate Hmg2-GFP was performed in wild-type (WT), *pah1Δ, cho2Δ, opi3Δ, ino4Δ,* and *spo7Δ* cells with chromosomally integrated Hmg2-GFP. (B) Quantification of (A). (**C**) Cycloheximide chase degradation assay for the ERAD-L substrate CPY*-3xHA was performed in WT and the indicated deletion mutants with CPY*-3xHA expressed from a centromeric plasmid. (D) Quantification of (C). (**E**) Cycloheximide chase degradation assay for the ERAD-C substrate Ste6-166-3xHA-GFP (Ste6*-HA-GFP)) was performed in WT and the indicated deletion mutants with Ste6*-HA-GFP expressed from a 2µ plasmid under the *PGK1* promoter. (F) Quantification of (E). For all assays, total protein levels were assessed using anti-Pgk1 blots as a loading control. Representative immunoblots from at least three independent biological replicates are shown. The means of the percentage of each ERAD substrate remaining from three biological replicates are plotted, with error bars representing the standard deviation (SD).

